# Mechanism of Dynamic Binding of Replication Protein A to ssDNA

**DOI:** 10.1101/2020.03.16.994681

**Authors:** Anupam Mondal, Arnab Bhattacherjee

**Affiliations:** School of Computational and Integrative Sciences, Jawaharlal Nehru University, New Delhi, India

**Author notes:** Author Contributions A.B. designed the research; A.M., and A.B. performed the research, analysed the data and wrote the paper.

**Keywords:** ssDNA-protein interactions, Replication protein A, dynamic binding

## Abstract

Replication protein A (RPA) serves as hub protein inside eukaryotic cells, where it coordinates crucial DNA metabolic processes and activates the DNA-damage response system. A characteristic feature of its action is to associate with ssDNA intermediates before handing over them to downstream proteins. The length of ssDNA intermediates differs for different pathways. This means RPA must have mechanisms for selective processing of ssDNA intermediates based on their length, the knowledge of which is fundamental to elucidate when and how DNA repair and replication processes are symphonized. By employing extensive molecular simulations, we investigated the mechanism of binding of RPA to ssDNA of different lengths. We show that the binding involves dynamic equilibrium with a stable intermediate, the population of which increases with the length of ssDNA. The vital underlying factors are decoded through collective variable principal component analysis. It suggests a differently orchestrated set of interactions that define the action of RPA based on the sizes of ssDNA intermediates. We further estimated the association kinetics and probed the diffusion mechanism of RPA to ssDNA. RPA diffuses on short ssDNA through progressive ‘bulge’ formation. With long ssDNA, we observed a conformational change in ssDNA coupled with its binding to RPA in a cooperative fashion. Our analysis explains how the ‘short-lived,’ long ssDNA intermediates are processed quickly in vivo. The study thus reveals the molecular basis of several recent experimental observations related to RPA binding to ssDNA and provides novel insights into the RPA functioning in DNA repair and replication.

**Significance Statement:** Despite ssDNA be the common intermediate to all pathways involving RPA, how does the latter function differently in the DNA processing events such as DNA repair, replication, and recombination just based on the length of ssDNA intermediates remains unknown. The major hindrance is the difficulty in capturing the transient interactions between the molecules. Even attempts to crystallize RPA complexes with 32nt and 62nt ssDNA have yielded a resolved structure of only 25nt ssDNA wrapped with RPA. Here, we used a state-of-the-art coarse-grained protein-ssDNA model to unravel the detailed mechanism of binding of RPA to ssDNA. Our study illustrates the molecular origin of variations in RPA action during various DNA processing events depending on the length of ssDNA intermediates.

## Introduction

Replication protein A (RPA) in eukaryotes plays a crucial role in DNA metabolic processes such as DNA replication, recombination, and repair (1–3). A common feature in all its pathways is that RPA interacts with ssDNA intermediates. For example, RPA rapidly coats the ssDNA regions during the initiation phase of DNA replication to provide stability and prevent the formation of secondary structures in ssDNA (4, 5). During the elongation phase, RPA coordinates the lagging strand polymerase switching and processing of Okazaki fragments (6). RPA also participates in DNA nucleotide excision repair (NER) process through its interactions with XPA to stabilize the open complex after locating the lesion and help in positioning the nuclease for the dual incision (7). Similarly, RPA sequesters the ssDNA during repair of DNA double-strand breaks (DSBs) and promotes the loading of RAD51 recombinase to form RAD51– ssDNA nucleoprotein filament (8).

Despite the myriad roles of RPA, the mechanism by which it can direct different ssDNA intermediates in distinct pathways to coordinate the various DNA metabolic processes remains obscure. The functions of RPA have been found to depend on its binding affinity to ssDNA and its ability to interact with other proteins(9) physically. Interestingly, RPA is a multi-domain protein, and its affinity for ssDNA is determined by a combined contributions from its six DNA binding domains (DBDs A-F). The DBDs are made of structurally-related, oligonucleotide binding (OB) folds, which are further grouped into three subunits, RPA70, RPA32, and RPA14 (see Fig 1). RPA70 is composed of four DBDs, namely F, A, B, and C (flexible F domain which remains disconnected from the rest of the complex is not shown in Fig 1A), whereas RPA32 and RPA14 comprise DBD-D with a winged-helix domain and DBD-E respectively. Among six DBDs, DBD-A and -B have the highest affinity (k_a_ ∼ 5×10^5^ M^-1^ and 5×10^4^ M^-1^respectively) for ssDNA (10, 11), whereas DBD-C and DBD-D exhibit weaker binding affinities (11–13). The latter two DBDs, along with DBD-E, form a trimerization core (14). The high affinity of DBD-A and DBD-B is primarily due to their interactions with ssDNA through a combination of polar and aromatic amino acids (15). This is evident from mutational analysis that suggests a significant reduction in affinity of RPA for ssDNA caused by mutations of polar amino acids. It is, however, interesting to note that the same is not valid for mutations at aromatic sites (11, 16). Mutations of aromatic residues have minimal impact on RPA affinity, but they are linked with a loss of DNA repair activity of RPA without any significant effects on its role in DNA replication(16). This indicates that while the affinity of RPA for ssDNA is essential for their assembly, the function of RPA in different pathways differ based on the details of intermolecular interactions. A difference also exists in the length of ssDNA intermediates involved in various DNA metabolic pathways (4, 11, 17, 18). For example, long (100-200 nt) but short-lived ssDNA intermediates are observed in DNA replication, whereas much shorter ssDNA intermediates (<30 nt) form a stable RPA-ssDNA complex for processing and repairing the DNA damage via the NER pathway (19). Does this mean the length of ssDNA intermediates regulate the action of RPA in various DNA metabolic pathways? If yes, what is the underlying molecular mechanism and what are the molecular determinants that differentiate the association of RPA to ssDNA based on the length of the latter? While answers to the questions are essential to know when and how specific DNA repair and recombination processes are orchestrated, very little has been explored so far.

**Figure 1.**
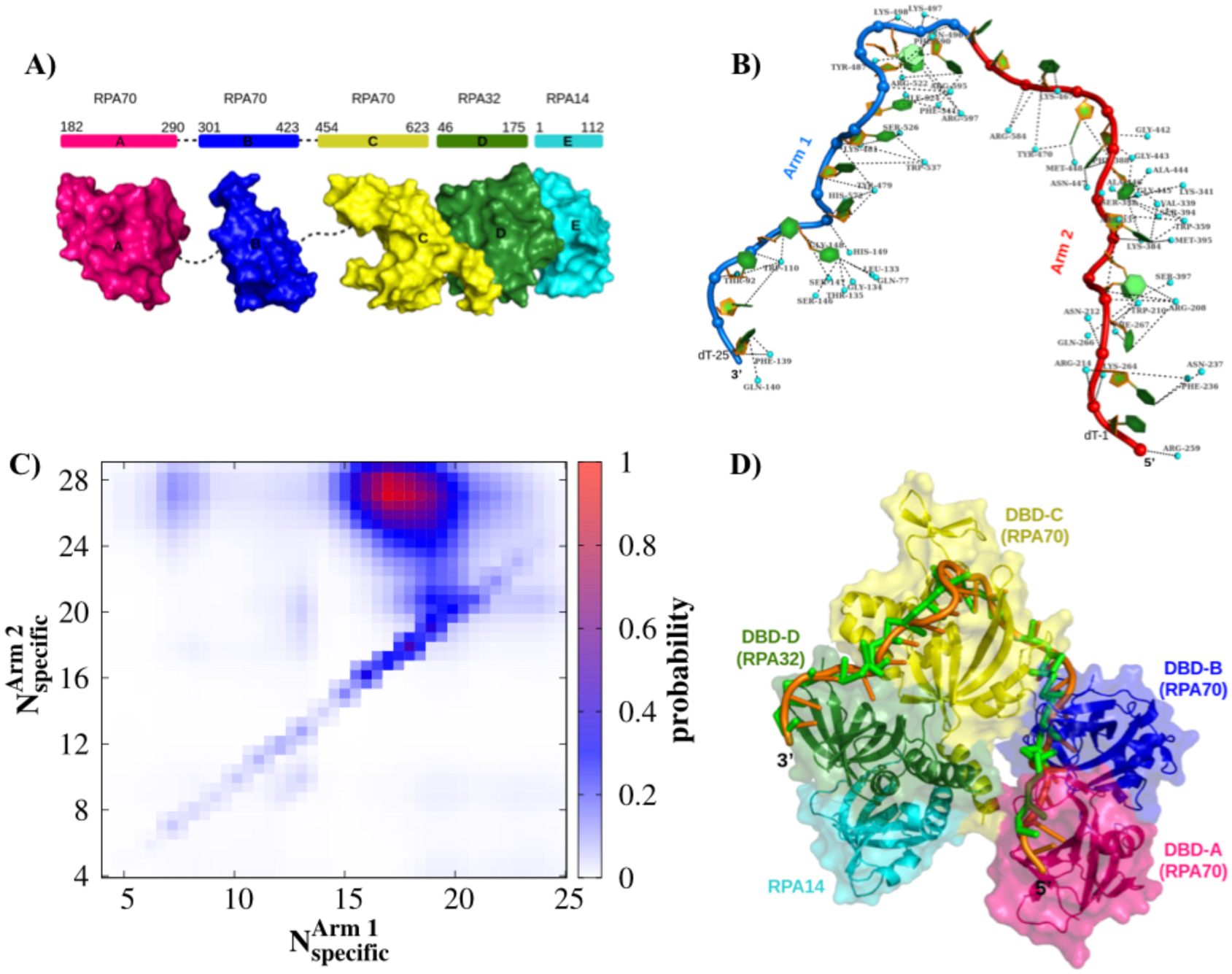
Structure and binding of RPA with ssDNA. (A) Schematic representation of the three RPA subunits RPA70, RPA32 and RPA14 with their residue numbers. RPA70 is composed of three DNA binding domains (DBD-A-C), whereas each RPA32 and RPA14 has DBD-D and DBD-E respectively. (B) The specific interfacial contacts of RPA residues (small cyan sphere) with ssDNA phosphate (blue sphere), sugar (orange colour) and base atoms (green ring). The ssDNA length is divided into approximately two equal halves and referred to as *arm-1* (13-25 nucleotides, blue colour) and *arm-2* (1-12 nucleotides, red colour). (C) Arm-wise specific interfacial contact formation probabilities are presented. The red regime indicates the bound state. (D) Crystal structure of the DBDs of *U. maydis* RPA bound to (dT)_25_ ssDNA (PDB ID 4GNX). The simulated ssDNA (green colour) is superimposed with the crystal one (orange color) after achieving the proper bound state conformation.

In this study, we have computationally explored the RPA-ssDNA binding energy landscape using a state-of-the-art coarse-grained (CG) protein-DNA model to investigate their association mechanism at molecular details. The model parameters were precisely tuned by employing the model in successfully predicting the binding of different ssDNA sequences to their respective protein partners. Our results suggest the presence of two distinct RPA-ssDNA binding modes that are connected via a dynamic equilibrium. The relative population of these states is a function of the length of ssDNA intermediates and their stability is governed by a set of intermolecular interactions. By applying a collective variable principal component analysis (PCA), we analysed the weights of each component of intermolecular interactions. The finding, together with the mechanistic details of ssDNA propagation on RPA, unravels an unanticipated mechanism of their association and provides crucial insights into the RPA functioning in DNA repair and replication.

## Results

### Binding of RPA to ssDNA varies with the length of ssDNA

To begin with, we first test if our model has achieved the correct balance between the weighted strengths of various protein-nucleic acid interactions and the conformational flexibility of the ssDNA tract to capture the RPA-ssDNA binding dynamics. For this, we performed binding simulations of a (dT)_25_ ssDNA with a RPA molecule starting from a completely unbound state. To monitor the progress of RPA association to ssDNA, we select the specific interfacial contacts between them as a reaction coordinate. An interfacial contact is recognised if two non-hydrogenous atoms, each from an amino acid of RPA and a nucleotide of ssDNA are within 3.7 Å in the crystal structure of RPA-(dT)_25_ (PDB ID: 4gop.pdb) complex. We found 57 such contacts (17 charged, 13 aromatic, 15 polar and 12 hydrophobic residues). The analysis is presented in Fig 1B, which closely resembles the RPA-ssDNA contacts probed by Fan et. al. (20). The heterogeneity in the type of interfacial residues suggests an orchestrated set of interactions behind the formation of the bound state complex. Domain wise, these contacts are spread over all four DBDs (A-D), among which DBD-C forms the maximum number of specific interfacial contacts with ssDNA despite its lowest affinity towards ssDNA. Towards this end, we note that the DBDs of RPA are interconnected by flexible linkers that allow them to interact independently with the ssDNA and assume an ensemble of conformations. Therefore, to understand if the specific interfacial contacts are formed sequentially along the length of ssDNA, we divided the length of the ssDNA into approximately two halves. The strand containing consecutive 13 nucleotides from the 3’-end is referred as *arm-1* and feature ∼27 specific contacts 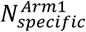, 8 charged, 7 aromatic, 8 polar and 4 hydrophobic residues), whereas 12 nucleotides from the 5’-end is termed as *arm-2* that contains ∼30 specific contacts 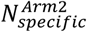, 9 charged, 6 aromatic, 7 polar and 8 hydrophobic residues). We estimated the arm-wise specific contact formation probabilities from trajectories of RPA-(dT)_25_ binding simulations. The result presented in Fig 1C shows the most probable region corresponds to a high number of specific contacts in both *arm-1* and *arm-2* of ssDNA. This essentially indicates capturing the correct bound state of RPA-(dT)_25_ in our simulations, which is also supported by an excellent structural overlap (see Fig 1D) obtained from the superimposition of our simulation generated bound state of RPA-(dT)_25_ on its crystal structure. Fig 1C also suggests that the route to the bound state of RPA-(dT)_25_ goes through a simultaneous formation of the specific interfacial contacts (evident from the diagonal data set) along both the arms of ssDNA. This means binding of different DBDs of RPA to ssDNA is not sequential, rather coherent.

Having seen that our model precisely captures the heterogeneity in RPA-ssDNA binding, we turned to investigate how RPA interacts with ssDNA intermediates of different lengths. It is important to note that previously complexes of RPA with a short (32 nt) and a long (62 nt) ssDNA molecule were crystallised (4gop.pdb and 4gnx.pdb) with 3.1 Å and 2.8 Å resolutions respectively. Interestingly, only 25 nt long identical ssDNA structures were resolved in both the complexes. While this suggests that ∼25 nt ssDNA stretch is sufficient to cover all four DBDs (A-D) of RPA, it fails to capture all the dynamic and transient contacts between nucleobases of ssDNA and amino acid residues of RPA that might define the differences in functioning of RPA during DNA replication and repair. Therefore, we probed the dynamics of binding of RPA separately with (dT)_25_ (short ssDNA), (dT)_32_ (dT)_40_ (dT)_50_ and (dT)_62_ (long ssDNA). The free energy profiles corresponding to short ((dT)_25_) and long ((dT)_62_) ssDNA with respect to the number of arm-wise contacts between nucleobases of ssDNA and amino acid residues 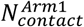 and 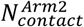 of RPA are presented in Fig 2A-B. The same for the other lengths of ssDNA can be found in Fig S3 in Supplementary text. Note that unlike 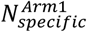 and 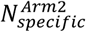, here we consider the formation of contacts whenever a nucleobase of ssDNA comes within 6 Å to an amino acid residue of RPA in our CG model. This allows us to monitor the progress of RPA binding to ssDNA even when they are non-specifically interacting. Our results show distinctly different free energy surfaces for (dT)_25_ and (dT)_62_. A single basin (denoted by H_25_) is observed in the free energy surface of RPA-(dT)_25_. The cluster of snapshots in Fig 2C corresponding to the basin H_25_ shows a horse-shoe shaped binding mode similar to its crystal structure (4gop.pdb). In contrast, the free energy surface of RPA-(dT)_62_ shows two basins (L_62_ and H_62_). L_62_ represents a stable intermediate state in RPA-(dT)_62_ binding where 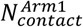 is significantly high (∼28 contacts) but 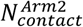 is limited to very few contacts (∼5 – 10 contacts). A cluster of snapshots (Fig 2D) corresponding to this state (state L_62_), show almost a linear conformation adopted by ssDNA on the RPA surface. In these conformations, the *arm-1* of ssDNA is associated with DBD-D and DBD-C of RPA, but *arm-2* remains flanking on RPA surface. The second basin (H_62_) on the free energy surface of RPA-(dT)_62_ corresponds to high 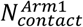 and 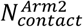 (∼28 contacts on each arm). The cluster of snapshots corresponding to H_62_ presented in Fig 2D elucidates a conformational transition from linear ssDNA of L_62_ to a ‘horse-shoe’ shaped bent conformation such that it can bind to DBD-B and DBD-A of RPA. We estimated the associated change in free energy, which suggests state-H_62_ is slightly more stable (ΔG ∼ 0.7 ± 0.001 kcal/mol, see Fig S4) compared to L_62_. The small free energy gap signifies a dynamic interconversion equilibrium exists between states L_62_ and H_62_. An analogous free energy gap (∼1.2 k_B_T) was reported for protein diffusion around the major grooves of dsDNA during its ‘sliding dynamics’ (21). The variation in the dynamic binding of RPA to ssDNA depending on the length of the latter is further manifested from our binding studies of RPA with an intermediate length of ssDNA molecules ((dT)_32_, (dT)_40_, (dT)_50_). Fig S5 shows as the length of ssDNA intermediate increases, the population of state-L_n_ (n is length of DNA) rises, indicating a shift in binding mechanism of RPA with the change in length of ssDNA intermediates. To this end, we also compared the stability of bound states of RPA with (dT)_25_ (state H_25_) and (dT)_62_ (state H_62_). We find the former is more stable (ΔG ∼ 0.46 ± 0.001 kcal/mol), which is in agreement with the observation that RPA forms a more stable complex with short ssDNA intermediate during DNA repair compared to the long ssDNA tract in DNA replication (19).

**Figure 2.**
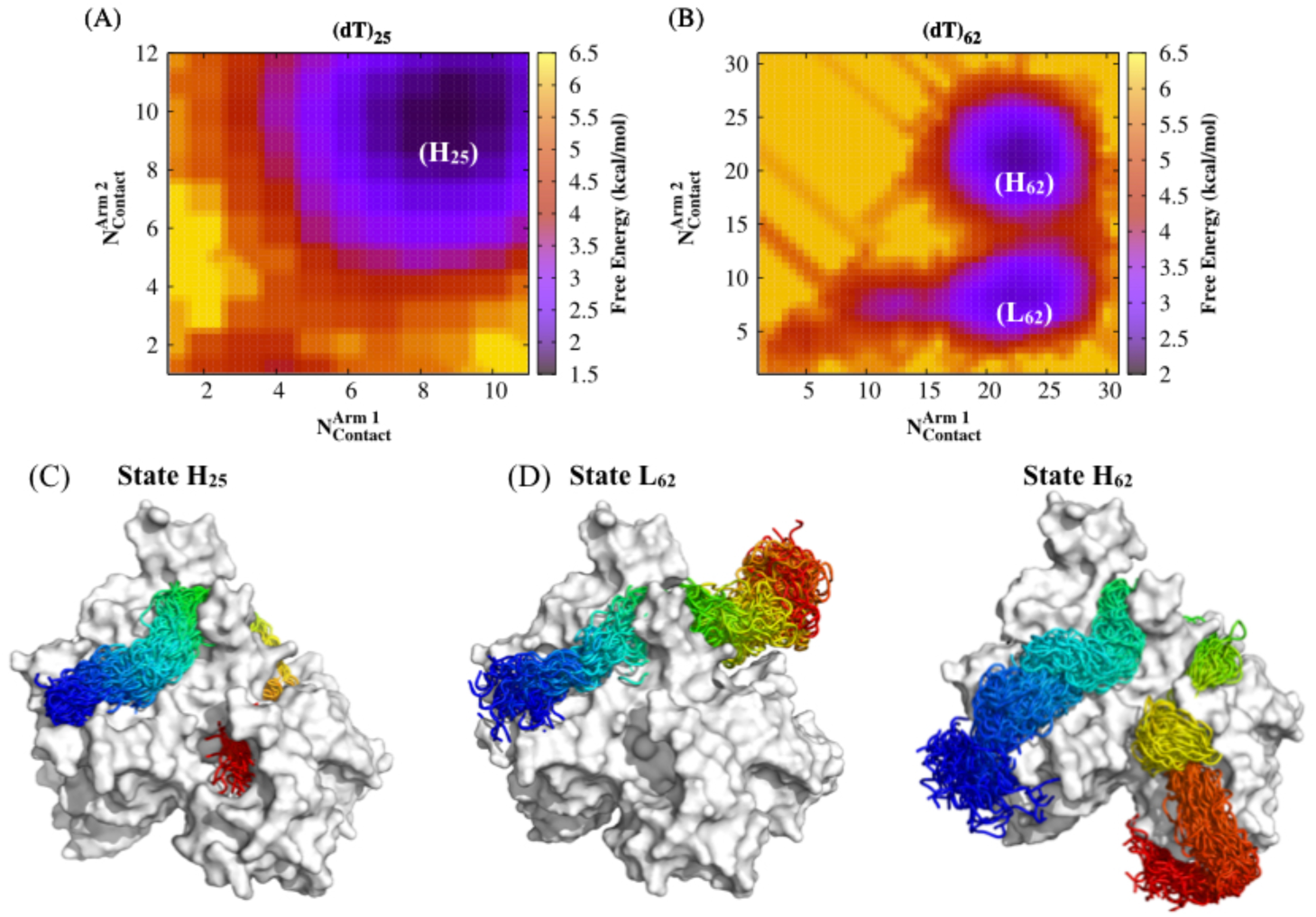
Length dependency in RPA-ssDNA binding. (A-B) Two dimensional free energy profiles for short ((dT)_25_) and long ((dT)_62_) length of ssDNA as a function of arm-wise contacts 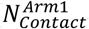 and 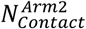 between nucleobases and the closest amino acid residues (within a distance cutoff of 6 Å). The basin *H*_*25*_ in (dT)_25_ indicates horse-shoe shaped bound state, whereas the two minima *L*_*62*_ and *H*_*62*_ in (dT)_62_ corresponds to a linear and a horse-shoe shaped conformation respectively. (C-D) Superposition of the cluster of snapshots corresponding to *H*_*25*_, *L*_*62*_ and *H*_*62*_ states are presented. RPA is shown in grey surface and ssDNA in rainbow colour.

### Molecular determinants of dynamic binding of RPA to ssDNA

What could be the physical basis of differences in RPA binding mechanism to ssDNA of different lengths? To understand this, we employed a collective variable (CV) principal component analyses (PCA) (22) that can pull apart the hidden cross-correlations among the collective variables. Thus, the technique is advantageous over the free energy analysis based on projection solely onto a single input variable. Here, as input basis variables, we considered different types of interactions between the amino acid residues of RPA and nucleotides of ssDNA that work in symphony to lead to the formation of RPA-ssDNA bound complex. This includes the following collective variables (CV1 to CV4): (1) electrostatic energy (E_elec_(RPA:ssDNA)) between phosphate residues of ssDNA and charged amino acids of RPA, (2) aromatic interaction energy (E_aro_), (3) hydrophobic interaction energy (E_hp_) (4) polar interaction energy (E_polar_) between the nucleobases of ssDNA and the relevant classes of amino acids of RPA. Our aim is to re-analyse the binding free energy profiles of RPA with (dT)_25_ and (dT)_62_.

Fig 3 and 4 present the free-energy surfaces of RPA with (dT)_25_ and (dT)_62_ respectively plotted against various estimated principal components (PCs). There are four principal components available (PC1 to PC4) that represent a new basis set of the transformed coordinates. The basis sets are obtained from linear combinations of the collective basis variables that we have taken as input to PCA. The coefficients used in the linear combinations and the variance are listed in Table 1 and 2 for (dT)_25_ and (dT)_62_, respectively. For moderate lengths of ssDNA, analysis were done and presented in supplementary text (Fig S6, S7 and Table S12 and S13). Our analysis suggests that E_elec_(RPA:ssDNA) is the major driving force (common to all PCs) for forming the bound state complex. We, therefore, consider E_elec_(RPA:ssDNA) as a common coordinate and presented the free energy profiles of the systems individually against each principal component. For example, the first panel of Fig 3 and 4 represent the free energy surfaces projected onto the first principal component (PC1) that account for ∼43% and ∼42% of the total variance. In addition, PC2 and PC3 are also important components that together with PC1 present 95.3% and 96% of the total variances in RPA-(dT)_25_ and RPA-(dT)_62_ assembly respectively. The variation of PC4 is less than 5%, indicating its minimal impact in regulating the RPA-ssDNA assembly. A comparison of the associated free energy surfaces of the two systems confirms the presence of a single basin for RPA binding to short ssDNA but two distinct basins for long ssDNA. The result suggests that the RPA-(dT)_62_ complex undergoes a dynamic interconversion equilibrium between the binding modes corresponding to the two basins. For RPA-(dT)_25_ the basin appears at E_elec_(RPA:ssDNA) = −25 kcal/mol, whereas the basins for RPA-(dT)_62_ complex appear at E_elec_(RPA:ssDNA) = −35 kcal/mol and −55 kcal/mol respectively. The increasing electrostatic energy with the increasing length of ssDNA is due to the involvement of more number of phosphate residues of ssDNA interacting with the positively charged amino acids of RPA. Another important insight emerges from the analysis of the combinations of collective basis variables corresponding to the significant principal components. In RPA-(dT)_25_ binding, PC1, PC2 and PC3 are made up of approximately 27%, 61% and 12% E_polar_. Together these account for ∼39% polar interactions between the nucleobases and the polar amino acids of RPA that stabilize the bound RPA-(dT)_25_ complex. The analysis explains why mutations of the polar residues in RPA lower its affinity for short ssDNA (11, 16). Similar estimations for E_aro_ and E_hp_ indicate that the contributions of interactions of nucleobases with aromatic and hydrophobic amino acids towards the precise binding of RPA to (dT)_25_ are 9.3% and 23.1% respectively. In contrast, the total contributions from polar, hydrophobic and aromatic interactions in the first three PCs for the RPA-(dT)_62_ binding are 13.2%, 34.13% and 27.6% respectively. The comparable weights of all three types of interactions suggest that more heterogenous intermolecular interactions drive the binding of RPA to long ssDNA. The interactions work in symphony to regulate the binding process and reduction of one component due to point mutations of a particular type of amino acids can be easily compensated by the other interactions. This is supported by the experimental observation that mutations of aromatic residues in RPA only minimally impact its affinity for ssDNA (16). Apart from polar and aromatic interactions, we found hydrophobic amino acids also play a significant role. We note that mutations of hydrophobic residues such as F238A, F269A, and F386A have reduced the affinity of RPA to ssDNA (16). However, it is difficult to assess to what extent the impact is due to a reduction in hydrophobic interactions as the mutations cause a simultaneous loss in aromatic interactions as well. The differences in the interaction pattern of RPA with varying length of ssDNA, thus define its differences in the mechanism of dynamic binding to ssDNA during DNA repair and replication processes.

**Table 1.** Coefficients for Principal Components (PCs) for (dT)_25_ binding mode

| | $E_{elec}$ | $E_{aro}$ | $E_{hp}$ | $E_{polar}$ |
| --- | --- | --- | --- | --- |
| PC1 (43.5%) | 0.295 | 0.037 | 0.394 | 0.273 |
| PC2 (38.9%) | 0.304 | 0.036 | 0.053 | 0.607 |
| PC3 (12.9%) | 0.204 | 0.456 | 0.221 | 0.120 |
| PC4 (4.7%) | 0.198 | 0.470 | 0.332 | 0 |

**Table 2.**
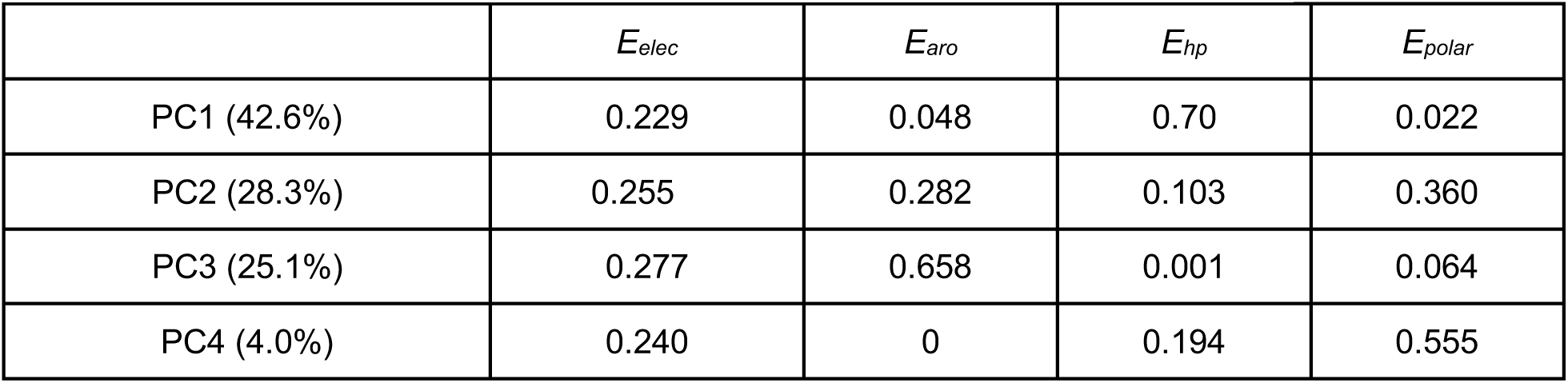
Coefficients for Principal Components (PCs) for (dT)_62_ binding mode

| | $E_{elec}$ | $E_{aro}$ | $E_{hp}$ | $E_{polar}$ |
| --- | --- | --- | --- | --- |
| PC1 (42.6%) | 0.229 | 0.048 | 0.70 | 0.022 |
| PC2 (28.3%) | 0.255 | 0.282 | 0.103 | 0.360 |
| PC3 (25.1%) | 0.277 | 0.658 | 0.001 | 0.064 |
| PC4 (4.0%) | 0.240 | 0 | 0.194 | 0.555 |

**Figure 3.**
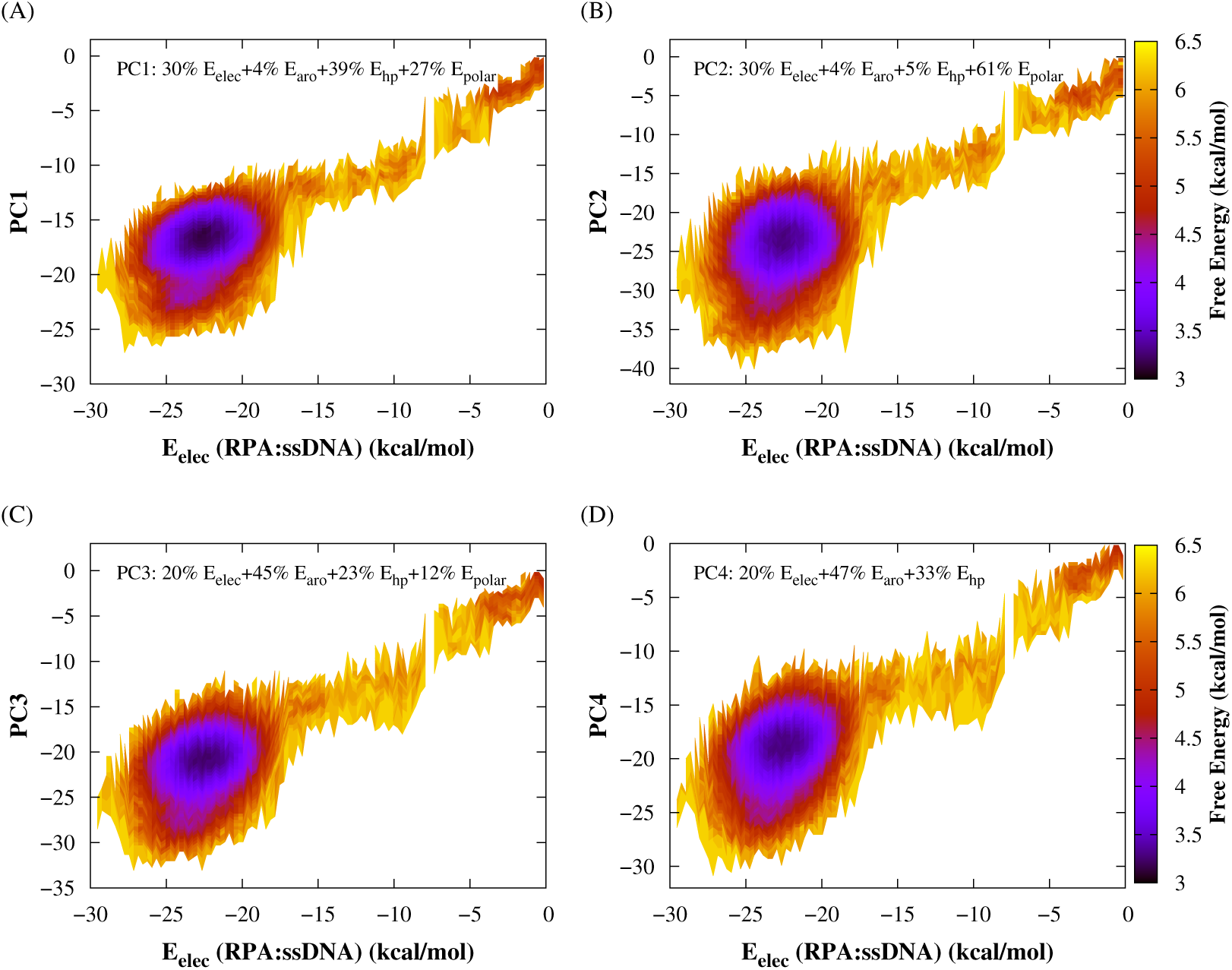
Two dimensional free energy surfaces for binding of RPA with (dT)_25_ as a function of various principal components (PCs). Each panel (A-D) shows the free energy is projected on each individual PC (PC1 to PC4) along with a common coordinate E_elec_(RPA:ssDNA). Each PC is a linear combination of the four collective basis variables in which the coefficients (number percentage) represents the fraction of these basis variables (see Table 1 for details).

**Figure 4.**
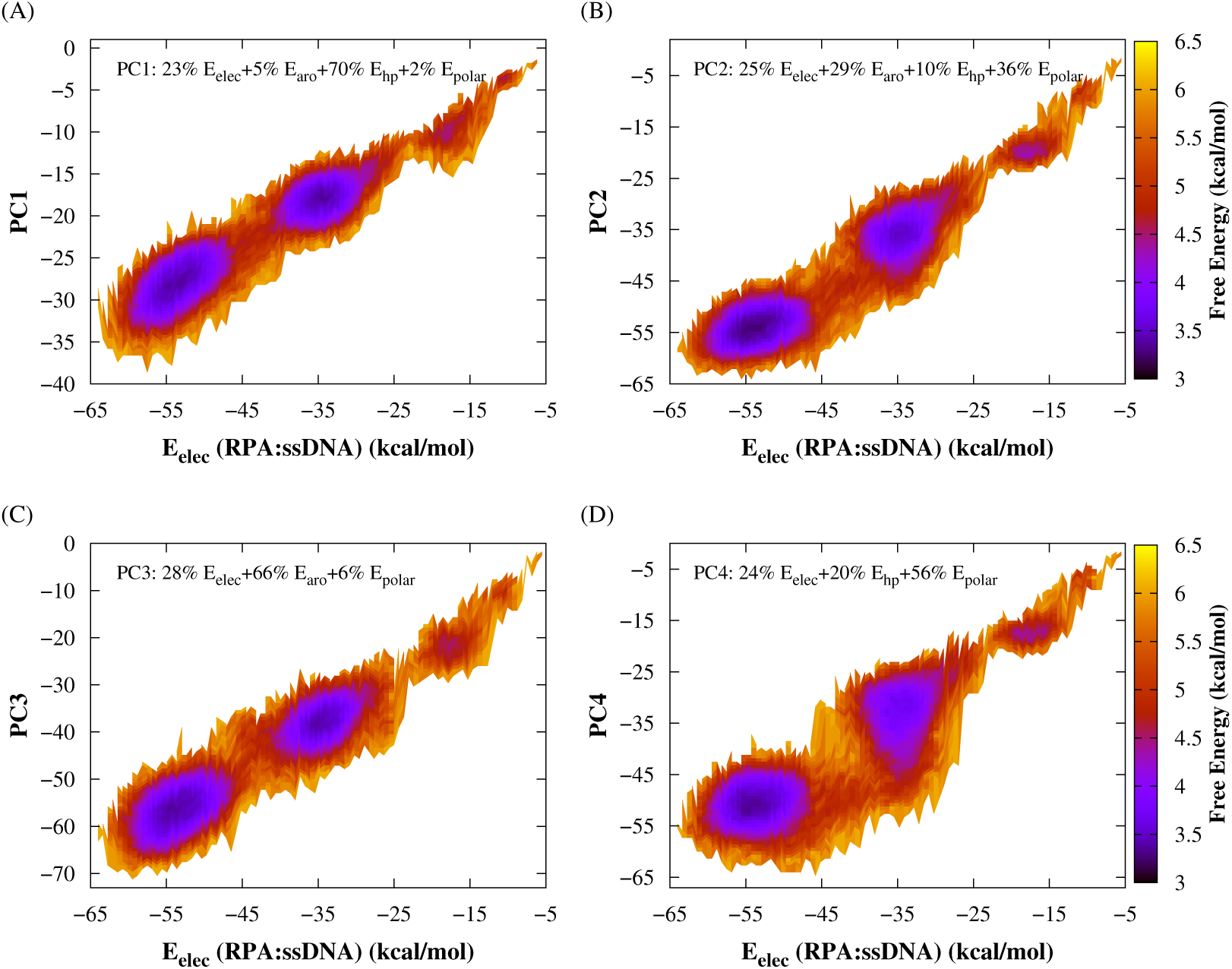
Two dimensional free energy surfaces for binding of RPA with (dT)_62_ as a function of various principal components (PCs). Each panel (A-D) shows the binding free energy is projected on a particular PC (PC1 to PC4) along with a common coordinate E_elec_(RPA:ssDNA). The number percentage in each PC represents the fraction of the collective basis variables (see Table 2 for details).

### Association mechanism of RPA differ for short and long ssDNA intermediates

Here, we investigate the molecular mechanism of RPA association to ssDNA, which requires probing the ssDNA propagation on RPA until it wraps the four DBDs (A-D) completely to form the bound state complex. Intuitively this is a long time-scale problem. Previous studies have rather investigated the translocation of ssDNA on protein in bound states only(23, 24). The analyses based on crystal structures of ssDNA bound E. coli SSB and RPA proteins demonstrate that short ssDNA segments(1-7nt) erect over the protein surface and form bulges due to defects in interactions between nucleobases of ssDNA and interfacial protein residues. With the dissolution of bulges, released nucleotides slide on the protein surface. The mechanism, referred as ‘reptation dynamics’, has been captured experimentally(25) although the formation of bulges is not. The picture is reminiscent of sliding mechanism of dsDNA on nucleosome via twist defect propagation(26).

To illustrate the RPA-ssDNA association mechanism, we start monitoring the movement of each nucleotide on RPA surface starting from when they established the first nonspecific contact with RPA. We noticed the formation of bulges on ssDNA as its characteristic feature. The bulges on RPA surface are identified if the distance (r_ij_) between i-th and j-th phosphate atoms of ssDNA is less than at least by 60% to its Kuhn length in a stretched conformation and the first phosphate residue participating in a bulge is positioned within 10 Å from the RPA surface. Fig 5A and B represent characterization of such bulges and suggest the most preferred size of the bulges are 3-4 nt long and they are transient (lifetime decays exponentially, see Fig 5B). Too small or large bulges are energetically unstable. The result is in agreement with the previous studies done on ssDNA and RPA protein(23, 24). To probe if the formation of bulges on ssDNA helps it to associate with RPA, we studied the time evolution of all phosphate residues that appear first when a bulge forms. Green dots present indices of those phosphate residues in Fig 5C, D, and E representing the movement of bulges in (dT)_25_ and arm-1 and arm-2 of (dT)_62_ respectively. The results reveal three intriguing aspects of bulge propagation in the context of the association mechanism of RPA to ssDNA: (i) bulges form throughout the length of (dT)_25_ tract. Some of them dissolute quickly and do not propagate with time. These are ‘static bulges’ observed previously for ssDNA sliding on E. coli SSB protein. The net movement of ssDNA due to the dissolution of such bulges is modest. Alternatively, some bulges are dynamic that aid in transporting nucleotides of ssDNA on RPA surface. (ii) The dynamic bulges move from 3’ to 5’ direction in (dT)_25_ ssDNA. Snapshots in Fig 5F taken at different time steps clearly portray the propagation of the bulges along the DBDs of RPA, leading to formation of the final RPA-(dT)_25_ association complex. The molecular driving force for the directed motion of the bulges is possibly the long-ranged electrostatic interactions. (iii) A similar directed movement of bulges can also be noticed in arm-1 of (dT)_62_. However, unlike arm-1 of (dT)_62_, a significant length of *arm-2* does not show any bulge formation (a black region in Fig 5E) up to MD steps ∼ 12 × 10^6^ MD. This corresponds to state L_62_ (see Fig 2), where ssDNA assumes a linear configuration with approximately one-third of its length flanking on the RPA surface. Transition from this state to the final bound state (H_62_) may be accompanied by formation and dissolution of bulges (‘reptation dynamics’) as it is in RPA-(dT)_25_. However, our analysis of bulges in *arm-2* indicates they are mostly static. Therefore, we hypothesize that alternative association mechanism of RPA might exist that can annotate how the long ‘short-lived’ ssDNA intermediates (19) are rapidly processed by RPA during DNA replication.

**Figure 5.**
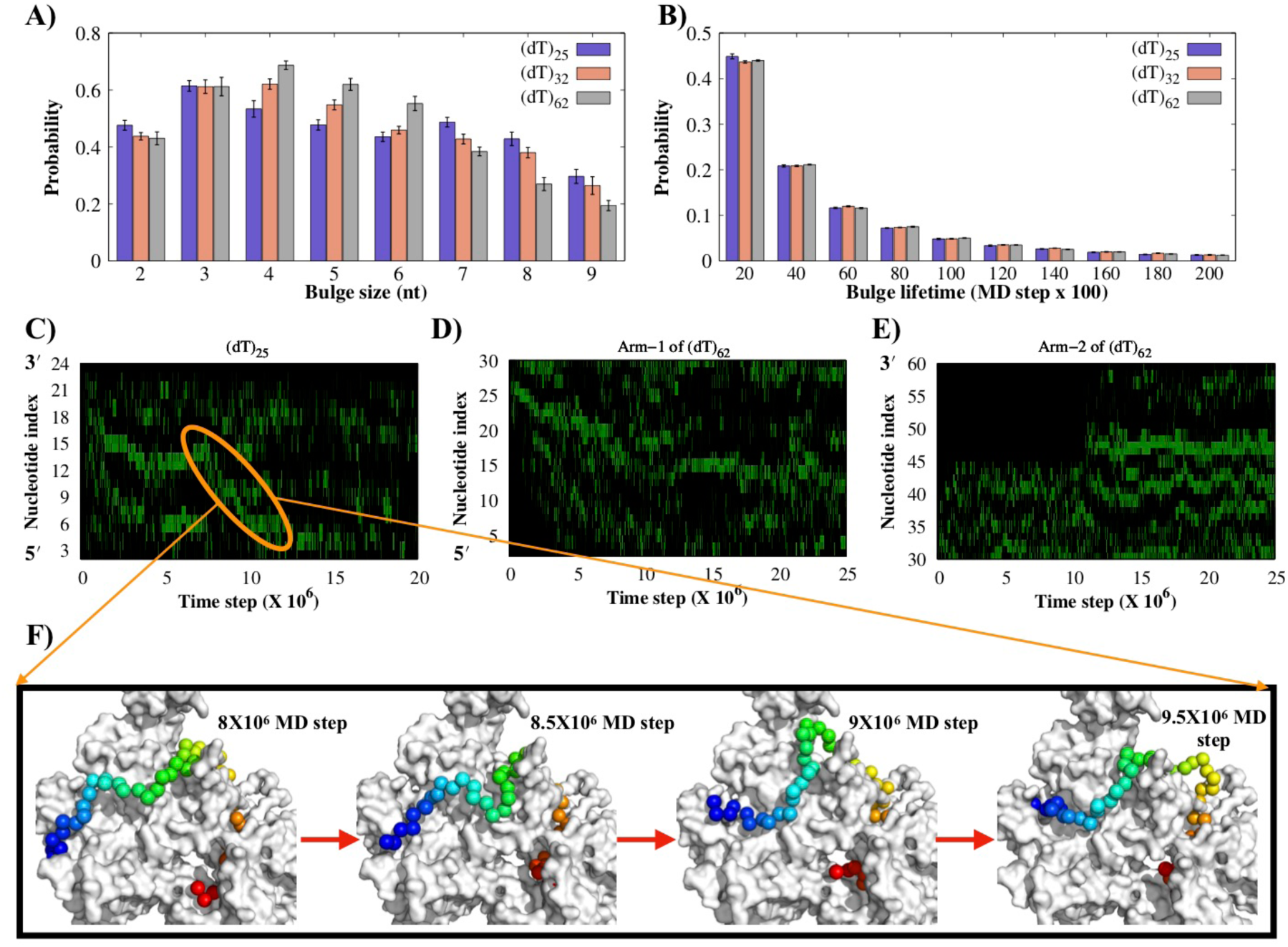
Characterization of ssDNA bulges on the RPA surface. (A-B) Probability distribution of different bulge sizes and their lifetime for (dT)_25_, (dT)_32_ and (dT)_62_ length of ssDNA. Bulges were quantified by examining the distance between those phosphate atoms which was less than at least 60% to its Kuhn length in a stretched conformation. (C-E) Time evolution of the nucleotide bulge index which was positioned on the RPA surface are shown for (dT)_25_ and *arm-1* and *arm-2* of (dT)_62_. 3’ and 5’ indicates the direction of the bulge movement. (F) Four snapshots, which were sampled at different MD time steps (highlighted region), represent the displacement of a bulge on the RPA surface. RPA is shown in grey surface and ssDNA in rainbow colour.

### Association kinetics and diffusion efficiency of RPA vary with the length of interacting ssDNA molecule

In order to test our hypothesis, we first estimated the association kinetics of RPA to ssDNA intermediates of different length. The rate of association was measured at DBD-A and DBD-D of RPA from the time required to form at least 80% of the specific RPA-ssDNA contacts. The choice of the DBDs is to ensure complete wrap of RPA by ssDNA as it does in the final bound complex. Our results presented in Fig 6A exhibits association rates of ssDNA at DBD-A and DBD-D respectively as a function of the length of ssDNA intermediates. We find there exist two different rates of ssDNA association to DBD-A for (dT)_25_, whereas a single for DBD-D. The observation is consistent with the experimental result based on interactions between surface tethered wild type RPA and Cy3-labeled (dT)_35_ that has measured two equilibrium constants (19) corresponding to a fast (K_a_ = 1.47 ± 0.27 × 10^9^ M^-1^) and a ten times slow binding (K_a_ = 1.66 ± 0.59 × 10^10^ M^-1^) mode of RPA to ssDNA. A recent study (9) has proceeded further to assess the individual contributions of different DBDs by site-specific labelling technique with MB543 and measured the association rate of (dT)_25_ at DBD-D given by k∼36.2 ± 2.3 s^-1^. In contrast, the association of ssDNA at DBD-A features a fast (k∼30.6 ± 9.8 s^-1^) and a slow kinetics (k ∼ 10.3 ± 9.8 s^-1^). The measurement of these rates from our simulations shows 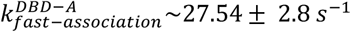 ∼27.54 ± 2.8 *s*^−1^ and 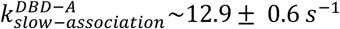 ∼12.9 ± 0.6 *s*^−1^, indicating an excellent match between simulations and experiment. However, our measured rate at DBD-D 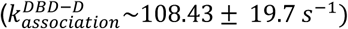 ∼108.43 ± 19.7 *s*^−1^) is ∼3 times higher than the experimental value. This could be because of the presence of winged helix associated with DBD-D, which we have not considered in the present simulations. Moving further, our analysis demonstrates that 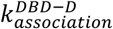 is roughly independent of the length of ssDNA intermediate. In comparison, 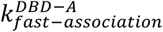 increases linearly (up to 3 times) with the increasing length of ssDNA intermediates. 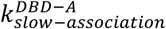 does not vary much with the length of ssDNA and even this binding mode does not exist for ssDNA length more than 40 nt. To explain the observations, we estimated the contact-time of specific interfacial RPA residues from the time they spent interacting (distance from nucleobase less than 6 Å) with at least one nucleotide. The analysis for both slow and fast binding events presented in Fig 6B clearly indicates a major role played by DBD-C and DBD-D of the trimerization core. The ssDNA spends significantly long time close to the interfacial residues of the trimerization core, resulting in a slow propagation of the ssDNA tract to DBD-A. This is counterintuitive as the DBDs of trimerization core has much lower affinity for ssDNA compared to DBD-B (k_a_ ∼ 5×10^4^ M^-1^) and DBD-A (k_a_ ∼ 5×10^5^ M^-1^). This is due to the high propensity of trimerization core to form more contacts with ssDNA that outcompetes the high affinity DBD-B and DBD-A. In some cases however, the conformational flexibility of ssDNA facilitates its rapid propagation without being stuck long at the trimerization core, resulting in a fast association kinetics at DBD-A. The impact of conformational flexibility is more prominent for longer ssDNA intermediates and it fully offsets the ability of trimerization core to retard the ssDNA association kinetics at DBD-A for ssDNA length greater than 40 nts. The slow binding mode thus does not exist for longer ssDNA. The same conformational feature (high conformational entropy) of long ssDNA also supports its extended conformation in state-L_n_ (n denotes the length of ssDNA, see Fig S5). The transition from state-L_n_ to the final bound complex of RPA-long ssDNA requires a conformational transition of ssDNA and its association at DBD-B and DBD-A. We note that both the processes happens cooperatively (see Fig S8 and Fig S9 in Supplementary text). Longer the ssDNA, greater is the cooperativity (see Fig S10 where cooperativity η_*H*_ increases linearly with ssDNA length). The molecular driving force is the long ranged electrostatic interactions between ssDNA and DBD-A and DBD-B. With the increasing flanking length of the flexible ssDNA, its capture radius increases that promotes faster nonspecific association of ssDNA at DBD-B and DBD-A(see Fig 6A, where 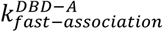 increases with ssDNA length). After initial nonspecific association, short ranged aromatic, polar and hydrophobic interactions act concertedly to form the final bound complex. The mechanism is analogous to the ‘fly-casting’ technique, where disordered proteins because of their high capture radius can associate faster to their binding partners (27–29).

**Figure 6.**
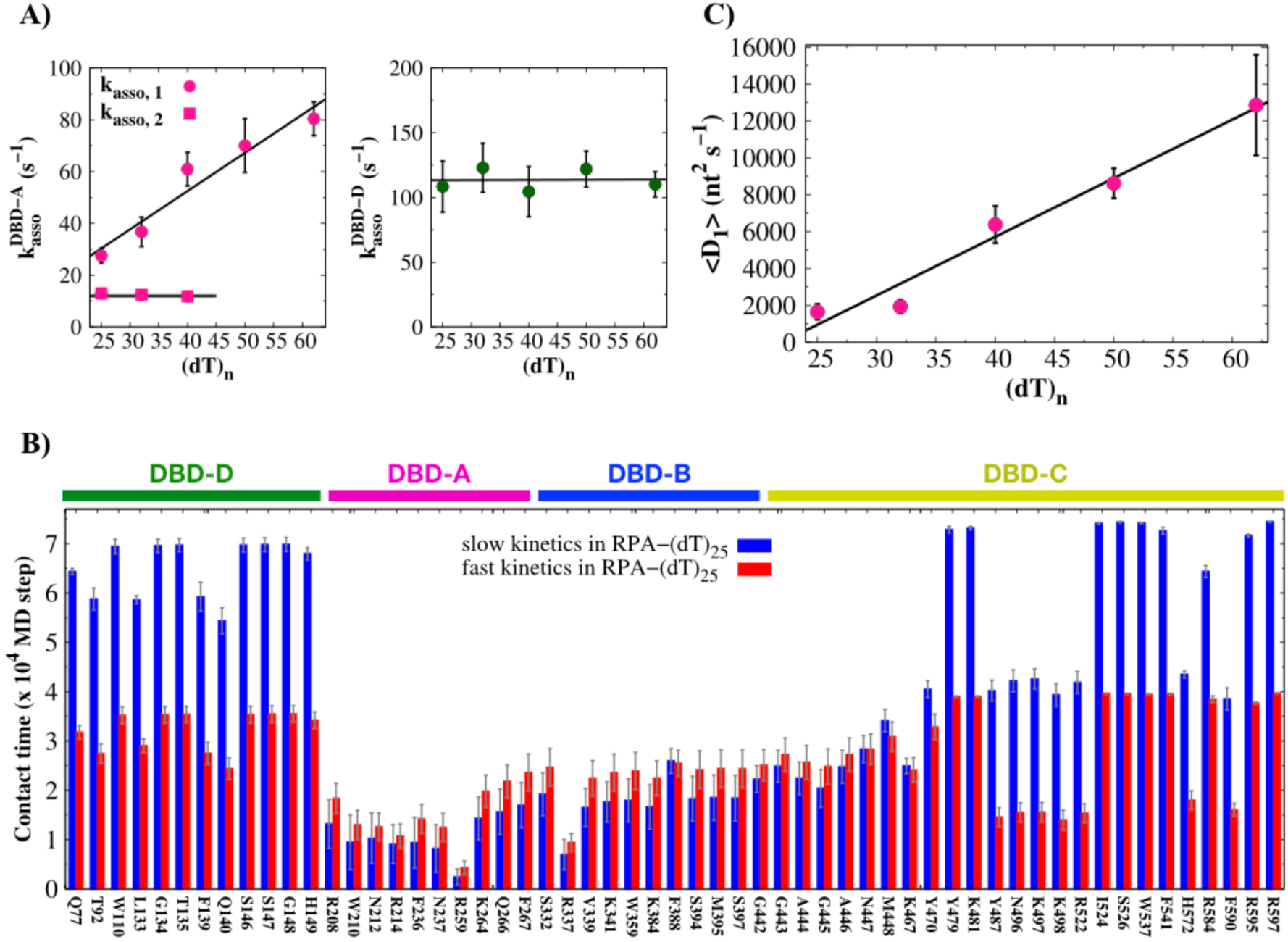
Kinetics of RPA binding with varying length of ssDNA. (A) The association rate of ssDNA to DBD-A and DBD-D of RPA with respect to ssDNA length (dT)_n_, where ‘*n*’ denotes the length of the ssDNA. The rates for 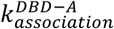 are best fit to two association kinetics, whereas the rate for 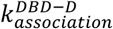 fit to one association kinetics. (B) The contact time of nucleotide bases in (dT)_25_ with interfacial RPA residues of different DBDs for slow and fast binding events. The range of each individual DBD is mentioned by a thick line at the top. (C) Variation in the 1D diffusion coefficient D_1_ at DBD-A of RPA as a function of (dT)_n_.

In addition to the association kinetics, we also investigated the diffusion efficiency of RPA on ssDNA in its bound state. Despite their high-affinity binding, RPA needs to be displaced or repositioned quickly for unhindered action of the downstream proteins involved in various metabolic pathways. This is common to many ssDNA binding proteins (30, 31). We measured the diffusion efficiency of RPA from its one-dimensional diffusion coefficient (D_1_) at 291 K from the slope of mean-square-displacement (MSD) of nucleotide bases at DBD-A. Our estimated diffusion coefficient D_1_ for (dT)_25_ is ∼ 0.00033 nt^2^/MD time step. This corresponds to D_1_ ∼ 1650 ± 426 nt^2^ s^-1^(see Fig S11 and Supplementary text for D_1_ calculation) considering each MD step in our simulations of 2 ps. The value is comparable to experimentally reported D_1_ is 5000 ± 400 nt^2^ s^-1^ for hRPA on ssDNA at 310 K (4). We further note that D_1_ increases linearly by 8 fold (D_1_ for (dT)_62_ ∼12859 ± 2727 nt^2^ s^-1^) as length of ssDNA changes from 25 nt to 62 nt, indicating a significant role of length of ssDNA intermediates in determining the fate of RPA-ssDNA bound complex during its post-processing. Our analysis suggests the molecular origin of faster diffusion of RPA on longer ssDNA is the formation of greater number of ‘dynamic bulges’ (see Fig S12 in Supplementary text) compared to that on shorter ssDNA due to the higher heterogeneity in intermolecular interactions in the former.

## Discussion

In this paper, we have computationally explored the full binding energy landscape of RPA with ssDNA of different lengths using a state-of-the-art coarse-grained protein-DNA model. The applicability of the model has been tested extensively by employing it in capturing the binding of different ssDNA sequences to their respective protein partners. Our motivation is to understand how the same RPA molecule results in different outcomes in DNA metabolic pathways such as DNA repair, recombination, and replication, depending on its interactions with ssDNA intermediates of varying lengths. Our study reveals the underlying molecular mechanism that provides crucial new insights into how and when DNA repair/replication processes are orchestrated. For example, we observed that RPA binds very differently to short ((dT)_25_) and long ((dT)_62_) ssDNA tracts. The short ssDNA transiently associates with RPA and then through ‘reptation dynamics’, where dynamic bulges on ssDNA form and dissolute continuously, ssDNA tract slowly wraps the four DBDs (A-D) of RPA and forms the RPA-ssDNA bound complex as shown by a single basin in the respective free energy landscape. The formation and movement of bulges from 3’ to 5’ end are triggered by mainly electrostatic interactions and defects in polar interactions respectively. The overall binding is coherent and dynamic, where the flexibly interconnected DBDs of RPA interact individually with (dT)_25_ through innumerable ways to stabilize the association of ssDNA finally into a horse-shoe shaped bent conformation on the RPA surface. The coherency further suggests that the ssDNA associates with the two extreme DBDs of RPA (A and D) roughly with comparable association kinetics, as was confirmed by the experimentally observed kinetic rates at these DBDs as well. While this is direct evidence that ssDNA binding to RPA is not sequential (ssDNA does not propagate along the sequence of DBDs in RPA), some association events were also reported that indicate ∼ 3 times slower association kinetics of ssDNA at DBD-A. Our analysis elucidates a major role played by the trimerization core of RPA in this slow kinetics. The high propensity of trimerization core to form the maximum number of contacts with (dT)_25_ outcompetes the high-affinity DBD-B and DBD-A in halting the ssDNA longer close to the trimerization core, resulting into slower association kinetics.

In comparison, RPA association to longer ssDNA ((dT)_62_) is quite different, although the initial association between (dT)_62_ and RPA occurs through the same ‘reptation dynamics’. The major difference between the free energy profiles of RPA-(dT)_25_ and RPA-(dT)_62_ is that the latter shows two basins, indicating the formation of final bound state via a stable intermediate state. The structural characterization of the intermediate state shows a linear conformation adopted by (dT)_62_, where roughly half of (dT)_62_ is bound to DBD-D and DBD-C but the remaining half at 5’ end flanks on the RPA surface. Our analysis suggests the high conformational flexibility (entropy) favours the extended conformation of (dT)_62_, the direct evidence of which comes from linear increment of the population of the stable intermediate state with the increasing length of ssDNA molecules. The high conformational entropy of longer ssDNA is also beneficial to offset the impact of trimerization core in retarding the association kinetics of ssDNA at DBD-A. In fact, the slow association kinetics of ssDNA at DBD-A diminishes to non-existence for ssDNA length longer than 40 nt. The transition from here to the final bound RPA-(dT)_62_ complex is unique for longer ssDNA only. To this end our results reveal two interesting aspects. (i) Firstly, RPA-(dT)_25_ complex is more stable (ΔG ∼ 0.46 ± 0.001 kcal/mol) than RPA-(dT)_62_ complex, which validates the experimental observation that RPA associates more strongly with short ssDNA intermediates during DNA repair compared to the long ssDNA tract in DNA replication (19). The molecular basis is the higher heterogeneity observed in the interactions between RPA and (dT)_62_. We show the association of RPA to long ssDNA is stabilized by a weighted combination of electrostatic, aromatic, polar and hydrophobic interactions. Reduction of one component via point mutations of a particular type of amino acids can be easily compensated by the other interactions. The prediction is supported by the experimental observation that mutations of aromatic residues in RPA only minimally impact its affinity for ssDNA. In contrast, electrostatic and polar interactions define the free energy profile of RPA to short ssDNA ((dT)_25_). This explains why mutations of the polar residues in RPA lower its affinity for short ssDNA (11, 16). (ii) Secondly, the intermediate state, exclusive to longer ssDNA, is only marginally less stable (ΔG ∼ 0.7 ± 0.001 kcal/mol) compared to the RPA-(dT)_62_ complex. The low energy barrier indicates a dynamic equilibrium between the states, unlike a single bound state for RPA with short ssDNA. During the transition from the intermediate state to the RPA-(dT)_62_ bound complex, the ssDNA undergoes a conformational change and binds to DBD-B and DBD-A of RPA. We find the conformational switch in ssDNA is facilitated by a cooperative binding at DBD-B and DBD-A of RPA. The cooperativity originates from a ‘fly-casting’ mechanism, where the negatively charged phosphates on the flanking ssDNA segment senses the positively charged residues of RPA from far to trigger their association. With the increasing flanking length of ssDNA, its capture radius increases that promote faster association. The mechanism illustrates how short-lived, long ssDNA intermediates rapidly associate with RPA during DNA replication.

To summarize, our study unravels the mechanism of dynamic binding of RPA to ssDNA. It explains the molecular origin of variations in RPA action during various DNA processing events depending on the length of ssDNA intermediates. Furthermore, the advantage of longer ssDNA is observed in faster diffusion of RPA on the ssDNA tract as well, indicating an ssDNA length-dependent fate of RPA-ssDNA bound complexes during their post-processing. The study also lays the foundation to analyse the impacts of mutation in RPA on its function and investigate how other RPA interacting proteins (RIPs) such as RAD52 that has lower a affinity for ssDNA displaces RPA from its complex with ssDNA during DNA metabolism.

## Materials and Methods

We developed a coarse-grained model to study ssDNA-protein interactions. The model resolution is similar to a previously developed model (32, 33) but varies significantly in terms of energetics. We adopted a coarse-grained representation of protein, where each amino acid is represented by a single bead placed at the respective C_α_ position (34). The energetics of the protein is described by a structure-based potential (35) (for details, see the Supporting text), which represents the funnel-like energy landscape for protein folding (35) and has been used extensively in studying protein-protein (36) and protein-nucleic acid interactions (37–43).

The resolution of the ssDNA molecule is three beads per nucleotide (phosphate, sugar and nitrogenous base), placed at their respective geometric centers. The energetics of the ssDNA molecule follows that of 3SPN.2 model of DNA by Hinckley et. al. (44). The model has successfully captured the structural features of ssDNA including precise predict the persistence length of ssDNA in agreement to the experimental measurements.

We have considered two components to describe the ssDNA-protein interactions namely, (i) long-ranged and (ii) short-ranged interactions. The long-ranged interaction includes electrostatic interactions between negatively charged phosphate beads of ssDNA and charged amino acids (R, K, D, E) residues of protein. For pH is <7, we have considered His residue to be positively charged (see Table S11). We assigned a unit positive charge to Lys, Arg and His and a unit negative charge on Glu and Asp. The phosphate bead was assigned a negative charge of 0.6 to take counterion condensation into account. We modelled the electrostatic interactions by Debye-Hückel potential that considers salt effect. It is noteworthy that previously Debye-Hückel potential has been applied successfully to understand crucial aspects of protein-nucleic acid interactions (37–43, 45, 46) despite its limited applicability for dilute solutions only. The effective strength of protein-DNA interaction is scaled by a factor of 1.67 to bring the local charge of phosphate beads back to -1, as described in their previous works (47).

The short-ranged interactions include aromatic stacking interactions between aromatic amino acids (F, W, H, Y) and the nucleobases of ssDNA, hydrogen bond interactions between non-aromatic amino acids (both polar and hydrophobic) and ssDNA bases and repulsive excluded volume interactions between protein and ssDNA residues. The aromatic stacking interactions are modelled by a Lennard-Jones potential, where reweighted sequence-dependent pairwise interaction strength parameters 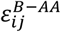 were chosen (48)(see Table S3). The reweighting factor is 0.6 to fit into our CG model in order to achieve the correct binding modes for various protein-ssDNA systems. The hydrogen bond interaction, similar to aromatic stacking, is also modelled by the Lennard-Jones potential, with the interaction strength of 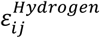 The non-aromatic amino acid residues (both polar and hydrophobic) have a strong tendency to form direct hydrogen bonds with nucleotides (20). We analysed the number of direct hydrogen bonds between non-aromatic amino acids and nucleobases from a set of 51 protein-ssDNA complexes (see Table S4). The profile shows a large variation in the interaction pattern between different protein residues with four different nucleotide sequences. We tuned the interaction strength values of 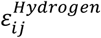 according to the hydrogen bond formation pattern analysed in Figure S1 and a complete list of sequence dependent pairwise base-non aromatic amino acid interaction strength values of 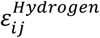 provided in Table S5. These values successfully capture the correct binding modes of various protein-ssDNA complexes (see Figure S2) irrespective of amino acid sequences of protein and nucleotide sequences of ssDNA molecule. The variety in sequence and structure of the studied complex strongly supports the generality and appropriateness of our model to study binding dynamics of ssDNA with proteins. Finally, a purely repulsive interaction is applied to consider the excluded volume effect. For further details, see the supporting information. To this end, it is important to note that our coarse-grained potential energy functions for the structure and energetics of the protein-ssDNA binding are fully transferable and does not include any bias towards the final specific complex.

The unbiased binding simulations were performed using Langevin dynamics with a friction coefficient *γ* = 0.1 kg/s. The integration time step used in the simulation is 0.05, which corresponds to a real time scale of 2 ps (49) (see Supporting material for time step conversion). The implicit solvent effect is considered by using the dielectric constant of water (78). Simulations were performed at temperatures as mentioned in the corresponding protein-ssDNA PDB complex (see Table S11), otherwise a temperature of 291K was used. The salt concentration is taken10 mM to ensure strong protein-ssDNA interactions. To explore RPA-ssDNA binding energy surface, we performed 50 independent simulations for each ssDNA length ((dT)_25_, (dT)_32_, (dT)_40_, (dT)_50_, and (dT)_62_). The simulations were 2 x 10^8^ MD steps long to ensure enough sampling of the conformational space.

## Acknowledgments

We gratefully acknowledge the financial support from DST India (DST PURSE, DST/INSPIRE/04/2013/000100, DST SERB CRG/2019/001001), and DBT CoE research grant. A.M. acknowledges financial support from CSIR India in the form of a Senior Research Fellow.

