## Supplementary text for "Mechanism of Dynamic Binding of Replication Protein A to ssDNA"

### This PDF file includes:

Supplementary text

Tables S1 to S13

Figures S1 to S12

References for SI reference citations

### Coarse-Grained Model for Proteins:

The protein is modelled through a coarse-grained (CG) representation where each protein residue is represented by one CG particle placed at the respective $C_\alpha$ position (1). The protein maintains its folded structure through a native topology-based model in which a Lennard-Jones potential encourages the formation of native contacts found in the crystal structure, thus help to preserve the protein native fold during the simulations (2). In addition, electrostatic interactions between positively charged (Arg, Lys, His) and negatively charged (Asp, Glu) amino acids are modelled through Debye-Hückel potential. We have considered His residues to be positively charged when the experimental pH value less than 7 was used for the crystallization of the protein-ssDNA complex deposited in the protein data bank (PBD) (see Table S11). Note that Debye-Hückel theory is inadequate to capture the effect of ion condensation and is valid only for dilute solutions. Despite several limitations, the Debye-Hückel potential has successfully been applied to understand many crucial aspects of nucleic acid biophysics (3–8).

The potential energy function for proteins is designated as:

$$E_{pot}^{Prot} = E_{bond}^{Prot} + E_{bend}^{Prot} + E_{torsion}^{Prot} + E_{Native-Contacts}^{Prot} + E_{ev}^{Prot} + E_{elec}^{Prot} \quad (1)$$

The bond energy, $E_{bond}^{Prot}$ between two successive $C_\alpha$ beads is given by,

$$E_{bond}^{Prot} = \sum_i k_b (r_i - r_i^0)^2 \quad (2)$$

where $k_b = 100\, kJ/mol/\AA^2$, $r_i$ and $r_i^0$ are the distances between $i^{th}$ and $(i+1)^{th}$ $C_\alpha$ beads in intermediate and folded structures of the protein respectively.

The bend energy, $E_{bend}^{Prot}$ for any variation in angle is given by,

$$E_{bend}^{Prot} = \sum_i k_\theta (\theta_i - \theta_i^0)^2 \quad (3)$$

where $k_\theta = 20\, kJ/mol/rad^2$, $\theta_i$ and $\theta_i^0$ are the angles between a vector connecting $i^{th}$ and $(i+1)^{th}$ $C_\alpha$ atoms and a vector connecting $(i+1)^{th}$ and $(i+2)^{th}$ $C_\alpha$ atoms in the intermediate and folded structures of the protein respectively.

The potential energy function for torsional angle, $E_{torsion}^{Prot}$ due to rotation in the dihedral angles between four consecutive $C_\alpha$ atoms connected by bonds is given as,

$$E_{torsion}^{Prot}=\sum_{i} k_{\varphi_1}[1-\cos 3(\varphi_i-\varphi_i^0)]+k_{\varphi_2}[1-\cos(\varphi_i-\varphi_i^0)] \quad (4)$$

where $k_{\varphi_1}=0.5\,kJ/mol$, $k_{\varphi_2}=1.0\,kJ/mol$, $\varphi_i$ and $\varphi_i^0$ are the torsional angles between $i^{th}$, $(i+1)^{th}$, $(i+2)^{th}$ and $(i+3)^{th}$ $C_\alpha$ beads in the intermediate and folded structures of protein respectively.

$E_{Native-Contacts}^{Prot}$ is the conformational energy that favours the formation of the native contacts found in the folded structure and such structure-based potential (as originally proposed by Clementi et. al. (2)) is modeled by a Lennard-Jones potential given by,

$$E_{Native-Contacts}^{Prot}=\sum_{i<j-3}^{native} \varepsilon_{ij}\left[5\left(\frac{\sigma_{ij}}{r_{ij}}\right)^{12}-6\left(\frac{\sigma_{ij}}{r_{ij}}\right)^{10}\right] \quad (5)$$

where $\varepsilon_{ij}=4.18\,kJ/mol$, $r_{ij}$ and $\sigma_{ij}$ are the distances between native pairs in the intermediate and folded structures of the protein respectively.

All the non-bonded and non-native $C_\alpha$ pairs are allowed to interact through a short-range repulsive excluded volume interactions, $E_{ev}^{Prot}$ given by,

$$E_{ev}^{Prot}=\sum_{i<j-3}^{non-native} \varepsilon_{ij}\left(\frac{\sigma_{ij}}{r_{ij}}\right)^{12} \quad (6)$$

where $\varepsilon_{ij}=1.0\,kJ/mol$, $r_{ij}$ denotes the distance between $i^{th}$ and $j^{th}$ $C_\alpha$ beads and $\sigma_{ij}$ is the sum of the radii of the interacting particles.

The Debye-Hückel screened electrostatic potential energy function is given by,

$$E_{elec}^{Prot}=\sum_{i<j}^{n_{elec}} \frac{q_i q_j e^{-r_{ij}/\lambda_D}}{4\pi\epsilon_0\epsilon(T,C)r_{ij}} \quad (7)$$

where $q_i$ and $q_j$ are the charges on $i^{th}$ and $j^{th}$ beads, $r_{ij}$ denotes the separation between them, $\lambda_D$ is the Debye screening length, $\epsilon_0$ is the dielectric permittivity of the vacuum and $\epsilon(T,C)$ is the dielectric permittivity of the solution.

The dielectric permittivity $\epsilon(T,C)$ is a function of salt molarity C and temperature T and can be expressed as the product of their individual contributions,

$$\epsilon(T,C)=\epsilon(T)a(C) \quad (8)$$

$$\epsilon(T)=249.4-0.788\,T/K+7.20\,X\,10^{-4}(T/K)^2 \quad (9)$$

$$a(C)=1.000-2.551\,C/M+5.151\,X\,10^{-2}(C/M)^2-6.889\,X\,10^{-3}(C/M)^3 \quad (10)$$

The Debye screening length can be written as $\lambda_D=\sqrt{\frac{\epsilon_0\epsilon(T,C)}{2\beta N_A e_c^2 I}}$ (11)

where $\beta=\frac{1}{k_B T}$ is the inverse thermal energy, $N_A$ is the Avogadro's number, $e_c$ is the elementary charge and $I$ is the ionic strength of the solution.

### Coarse-Grained Model for ssDNA:

In this study, we adopted 3SPN.2 coarse-grained model of DNA developed in de Pablo's group, where each nucleotide is described as three beads: one for the phosphate, one for the sugar and one for the base (9). The model accurately captures the correct persistence length for ssDNA which are in good agreement with experimental measurements, thus makes it a suitable candidate to study the DNA dynamics at the molecular level.

The potential energy function used to model the ssDNA is given by

$$E_{pot}^{ssDNA} = E_{bonded}^{ssDNA} + E_{non-bonded}^{ssDNA} \tag{12}$$

$E_{bonded}^{ssDNA}$ is a combination of bond, angle and torsional potentials to preserve the ssDNA initial structure, given by

$$E_{bonded}^{ssDNA} = E_{bond}^{ssDNA} + E_{bend}^{ssDNA} + E_{torsion}^{ssDNA}$$

$$= \sum_i k_b (r_i - r_i^0)^2 + 100 k_b (r_i - r_i^0)^4$$

$$+ \sum_i k_\theta (\theta_i - \theta_i^0)^2 + \sum_i -k_\varphi \exp\left(\frac{(\varphi_i - \varphi_i^0)^2}{2\sigma_{\varphi,i}^2}\right) \tag{13}$$

where $k_b = 0.6\, kJ/mol/Å^2$ and $r_i$, $r_i^0$ are the instantaneous and equilibrium bond lengths for the $i^{th}$ DNA bond respectively; $k_\theta = 200\, kJ/mol/rad^2$ and $\theta_i$, $\theta_i^0$ are the instantaneous and equilibrium bond angles for the $i^{th}$ bond angle of DNA respectively; $k_\varphi = 6.0\, kJ/mol$ and $\sigma_{\varphi,i}$, $\varphi_i^0$ are the Gaussian well depth and equilibrium angle for the $i^{th}$ dihedral respectively. In this model, the dihedral forces only act on the backbone of the system, i.e., the dihedrals are formed by phosphate and sugar sites (S-P-S-P and P-S-P-S).

For non-bonded potential, we use excluded volume interactions, electrostatic interactions and base-stacking interactions as given by,

$$E_{non-bonded}^{ssDNA} = E_{ev}^{ssDNA} + E_{elec}^{ssDNA} + E_{bstk}^{ssDNA} \tag{14}$$

The excluded volume interactions, $E_{ev}^{ssDNA}$ between sites i and j are modeled through a purely repulsive Lennard-Jones potential,

$$E_{ev}^{ssDNA} = \sum_{i<j} \begin{cases} \varepsilon_r \left[ \left( \frac{\sigma_{ij}}{r_{ij}} \right)^{12} - 2 \left( \frac{\sigma_{ij}}{r_{ij}} \right)^{6} \right] + \varepsilon_r & r < r_c \\ 0 & r \geq r_c \end{cases} \quad (15)$$

where $\varepsilon_r = 1.0\, kJ/mol$ and $r_c = \sigma_{ij}$ is the average diameter of the interacting particles. The potential only acts between those sites that are not involved in any bonded interactions or base stacking interactions.

The electrostatic interactions $E_{elec}^{ssDNA}$ between all charged phosphate atoms, which are not from neighbouring nucleotides, are modeled using Debye-Hückel potential energy function given in equation (6). The phosphates are assigned a negative charge of 0.6 instead of 1.0 in order to consider counter-ion condensation.

The base stacking interactions rely on a Morse potential of the form

$$U_{Morse}(\varepsilon_{ij}, \alpha_{ij}, r_{ij}) = \varepsilon_{ij} (1 - e^{(-\alpha_{ij}(r_{ij} - r_{ij}^0))})^2 - \varepsilon_{ij} \quad (16)$$

which is decomposed into a repulsive component

$$U_{Morse}^{rep}(\varepsilon_{ij}, \alpha_{ij}, r_{ij}) = \begin{cases} \varepsilon_{ij} \left( 1 - e^{\left( -\alpha_{ij} \left( r_{ij} - r_{ij}^0 \right) \right)} \right)^2 & r_{ij} < r_{ij}^0 \\ 0 & r_{ij} \geq r_{ij}^0 \end{cases} \quad (17)$$

and an attractive component

$$U_{Morse}^{attr}(\varepsilon_{ij}, \alpha_{ij}, r_{ij}) = \begin{cases} -\varepsilon_{ij} & r_{ij} < r_{ij}^0 \\ \varepsilon_{ij} \left( 1 - e^{\left( -\alpha_{ij} \left( r_{ij} - r_{ij}^0 \right) \right)} \right)^2 - \varepsilon_{ij} & r_{ij} \geq r_{ij}^0 \end{cases} \quad (18)$$

Here $\varepsilon_{ij}$ denotes the well depth of attraction between sites i and j, $\alpha_{ij}$ is used to control the range of attraction and $r_{ij}^{0}$ is the equilibrium distance between interacting sites.

A modulating function f is also incorporated to the angle of interaction in the base-stacking interaction, which is of the form

$$f(K, \Delta\theta) = \begin{cases} 1 & -\frac{\pi}{2K} < \Delta\theta < \frac{\pi}{2K} \\ 1 - \cos^2(K\Delta\theta) & -\frac{\pi}{K} < \Delta\theta < -\frac{\pi}{2K} \; or \; \frac{\pi}{2K} < \Delta\theta < \frac{\pi}{K} \\ 0 & \Delta\theta < -\frac{\pi}{K} \; or \; \Delta\theta > \frac{\pi}{K} \end{cases} \tag{19}$$

where the modulating constant K manages the attractive cone width. With these definitions, we can fully describe the potential energy function for the pi-stacking interactions in the form of base-stacking interactions as

$$E_{bstk}^{ssDNA} = \sum^{n_{bstk}} \begin{cases} U_{Morse}^{rep}\left(\varepsilon_{ij}, \alpha_{BS}, r_{ij}\right) + f\left(K_{BS}, \Delta\theta_{BS,ij}\right) U_{Morse}^{attr}\left(\varepsilon_{ij}, \alpha_{BS}, r_{ij}\right) & r_{ij} < r_{ij}^{0} \\ f\left(K_{BS}, \Delta\theta_{BS,ij}\right) U_{Morse}^{attr}\left(\varepsilon_{ij}, \alpha_{BS}, r_{ij}\right) & r_{ij} \geq r_{ij}^{0} \end{cases} \tag{20}$$

where $K_{BS}$=6.0, $\alpha_{BS}$=3.0, $\varepsilon_{ij}$ gives the depth of the well of attraction between interacting sites and $\theta_{BS}$ is the angle between the vector connecting sugar and base in the $5'$ direction and the vector joining the two base atoms in the $3'$ direction (see Table S1 and S2).

### Sequence-Dependent Model for Protein-ssDNA Interactions:

In our model, we incorporated the following four interaction potentials between protein and ssDNA molecules in order to study the binding of ssDNA with proteins at the molecular level: (i) the electrostatic interactions between negatively charged phosphate beads and charged amino acids (Arg, Lys, His, Asp, Glu), (ii) the aromatic stacking interactions between aromatic amino acids (Phe, Trp, His, Tyr) and the ssDNA bases, (iii) the hydrogen bond interactions between non-aromatic amino acids and ssDNA base beads and (iv) the repulsive excluded volume interactions between protein residues and other ssDNA beads. Therefore,

$$E_{pot}^{Prot-ssDNA} = E_{elec}^{Prot-ssDNA} + E_{Aromatic}^{Prot-ssDNA} + E_{Non-Aromatic}^{Prot-ssDNA} + E_{ev}^{Prot-ssDNA} \tag{21}$$

The electrostatic interactions between charged residues of protein and ssDNA are modelled using Debye-Hückel potential as given in equation (6). Since the phosphate beads are assigned a negative charge of 0.6 in the DNA model, the effective charge of interactions between protein and ssDNA sites is scaled by a factor of 1.67 in order to bring the local charge of phosphate beads back to -1, as used in the previous work (10).

Unlike dsDNA, the $\pi-\pi$ stacking interactions play an important role in protein-ssDNA interactions (11, 12). These stacking interactions between aromatic amino acid residues and the ssDNA bases are modelled by a Lennard-Jones potential of the form (13)

$$E_{Aromatic}^{Prot-ssDNA} = \sum \varepsilon_{ij}^{B-AA} [5\left(\frac{\sigma_{ij}}{r_{ij}}\right)^{12} - 6\left(\frac{\sigma_{ij}}{r_{ij}}\right)^{10}] \tag{22}$$

where $\varepsilon_{ij}^{B-AA}$ is the base-aromatic amino acid interaction strength, $\sigma_{ij}$ is the sum of the radii of the interacting beads, $r_{ij}$ is the distance between aromatic protein residue and DNA base in a given conformation during the simulation and a distance criterion of $10\mathring{A}$ is applied between them above which the inter-particle potential is zero. The interaction strength values of $\varepsilon_{ij}^{B-AA}$ varies differently depending on the sequence of ssDNA bases and the nucleobase-aromatic $C_\alpha$ bead pairs. We adopted the sequence-dependent pairwise base-aromatic interaction strength values from the experimental measurement of Rutledge et al. (11) (see Table S3). We reweighted the pairwise $\varepsilon_{ij}^{B-AA}$ values by a factor of 0.6 to fit into our CG model appropriately in order to achieve the correct binding modes for all studied protein-ssDNA systems.

In addition to stacking interactions between nucleobase and aromatic protein residues, the hydrogen bond interactions play a crucial role in stabilizing the interface between proteins and ssDNA (14). The non-aromatic amino acid residues (both polar and hydrophobic) have a strong tendency to form direct hydrogen bonds with nucleotides (15). We analyzed the number of direct hydrogen bonds (see Figure S1) from a set of 51 protein-ssDNA complexes (see Table S4). The profile shows a large variation in the interaction pattern between different protein residues with four different nucleotide sequences. Similar to $\pi-\pi$ stacking, we incorporated such hydrogen bond interactions between non-aromatic $C_\alpha$ atoms and the nucleotide base beads in the model through the same Lennard-Jones potential,

$$E_{Non-Aromatic}^{Prot-ssDNA} = \sum \varepsilon_{ij}^{Hydrogen} \left[ 5\left(\frac{\sigma_{ij}}{r_{ij}}\right)^{12} - 6\left(\frac{\sigma_{ij}}{r_{ij}}\right)^{10} \right] \tag{23}$$

The strength of the interactions $\varepsilon_{ij}^{Hydrogen}$ is tuned according to the interaction pattern analysed in Figure S1 and a complete list of sequence-dependent pairwise base-non aromatic interaction strength is provided in Table S5. These values successfully capture the correct binding modes of various protein-ssDNA complexes as compared to the crystal structures (see Figure S2).

Finally, a purely repulsive excluded volume interactions are applied to protein atoms and other ssDNA beads given by

$$E_{ev}^{Prot-ssDNA} = \sum_{i<j-3} \varepsilon_{ij} \left(\frac{\sigma_{ij}}{r_{ij}}\right)^{12} \tag{24}$$

where $\varepsilon_{ij} = 1.0\,kJ/mol$, $\sigma_{ij}$ is the average site diameter between i and j and $r_{ij}$ is the separation between them.

Thus, the total potential energy of the system becomes,

$$E_{pot} = E_{pot}^{Prot} + E_{pot}^{ssDNA} + E_{pot}^{Prot-ssDNA} \tag{25}$$

### Simulation Protocol:

Initially, we started the simulation by placing a free ssDNA (the length of which is same as that of the corresponding crystal structure) far away from the protein surface inside a simulation box with periodic boundary conditions. The binding of ssDNA to the proteins from the complete unbound state was studied using Langevin dynamics with a friction coefficient $\gamma = 0.1$ kg/s. The protein-ssDNA system was simulated using the dielectric constant of water (78) at a salt concentration of 10 mM. Note that due to coarse-grained nature of the model, the inter-bead electrostatic interaction is weaker compared to the all-atom model and thus, lower salt concentrations may correspond to the strong protein-DNA interactions compared (16) to the typically used ionic strengths in experiments. We used the temperature as mentioned in the corresponding protein-ssDNA PDB complex (see Table S11), otherwise, a temperature of 291K is used in this study. We performed 50 independent simulations for RPA-ssDNA system (10 for all other systems studied) in order to achieve the statistical robustness for each system. All simulations were $2\,x\,10^8$ MD steps long and performed in the canonical ensemble with Langevin Thermostat.

### Calibration of time scales in coarse grained MD simulations

In order to compare the results obtained from our coarse-grained (CG) simulations with experimental results, we mapped the time scale of our coarse-grained simulations with physical time units. For this conversion, we followed the prescription of Veitshans *et al (17)*.

A natural choice of the unit of time is $\tau_S = \left(\frac{m_0 a_0^2}{\epsilon_h}\right)^{1/2}$ (26)

where $m_0$ is the mass of the nucleobase, $a_0$ is the Van der Waals' radius of the nucleobase and $\epsilon_h$ is the average strength of the aromatic and non-aromatic interactions.

Since the ssDNA sequence is poly T for RPA-ssDNA system, therefore the values of $m_0$ and $a_0$ are $m_0 = 125.1\, amu = 2.08\, x\, 10^{-22}\, g$ and $a_0 = 3.55\, \mathring{A} = 3.55\, x\, 10^{-8}\, cm$. Also, the average value of the aromatic and non-aromatic interaction strengths is $\epsilon_h = 0.76\, kcal\, mol^{-1} = 5.32\, x\, 10^{-14}\, erg$.

Thus, we evaluate $\tau_S$ as : $\tau_S = \left(\frac{m_0 a_0^2}{\epsilon_h}\right)^{1/2} \simeq 2\, ps$.

From this mapping, we conclude that a single time step in our coarse-grained simulations is in the order of $\sim 2\, ps$.

The 1D diffusion coefficient $D_1$ for $(dT)_{25}$ obtained from our CG simulations is $\sim 0.00033\, nt^2 / MD\, time\, step$. Converting this time step into real time units (considering each MD step is equal to 2 ps), we obtain

$$D_1 = 0.00033\, nt^2 / MD\, time\, step = \frac{0.00033}{2 \times 10^{-12} \times 10^5} nt^2 s^{-1} = 1650\, nt^2 s^{-1}$$

[since $2\, ps = 2 \times 10^{-12} s$ and the factor $10^5$ in the denominator is the maximum time step up to which the MSD data behaves linearly (see Figure S11)].

**Table S1. The values of strength $\varepsilon_{ij}$ for base stacking interactions. ↑ and ↓ denotes the sense and anti-sense strands respectively. $3'$↑ and ↓$5'$ denotes adjacent bases in the $3'$ and $5'$ directions respectively.**

| | | Base $3'$↑ | | | |
|---|---|---|---|---|---|
| | | $\varepsilon$ $(kJ/mol)$ | | | |
| | | A | T | G | C |
| Base $5'$ ↑ | A | 14.39 | 14.34 | 13.25 | 14.51 |
| | T | 10.37 | 13.36 | 10.34 | 12.89 |
| | G | 14.81 | 15.57 | 14.93 | 15.39 |
| | C | 11.42 | 12.79 | 10.52 | 13.24 |

**Table S2. The values of equilibrium distances $\sigma_{ij}$ and equilibrium angles $\theta_{BSo}$ for base stacking interactions. ↑ and ↓ denotes the sense and anti-sense strands respectively. $3'$↑ and ↓$5'$ denotes adjacent bases in the $3'$ and $5'$ directions respectively.**

| | | Base $3'$ ↑ | | | | | | | | |
|---|---|---|---|---|---|---|---|---|---|---|
| | | $\sigma$ (Å) | | | | | $\theta_{BSo}$ (°) | | | |
| | | A | T | G | C | | A | T | G | C |
| Base $5'$↑ | A | 3.716 | 3.675 | 3.827 | 3.975 | A | 101.15 | 85.94 | 105.26 | 90.26 |
| | T | 4.238 | 3.984 | 4.416 | 4.468 | T | 101.59 | 89.50 | 104.31 | 90.82 |
| | G | 3.576 | 3.598 | 3.664 | 3.822 | G | 100.89 | 84.83 | 105.48 | 90.18 |
| | C | 3.859 | 3.586 | 4.030 | 3.957 | C | 115.95 | 101.51 | 119.32 | 104.49 |

**Table S3. The experimental values of strength $\varepsilon_{ij}^{B-AA}$ for specific $\pi-\pi$ stacking interactions between different nucleobase-aromatic bead pairs (as reported by Rutledge *et al.* (11)). The table values are adopted from the recent study on the binding of ssDNA and ssRNA with proteins** (18)**.**

| | A | T | G | C |
|---|---|---|---|---|
| PHE | 3.3 | 2.8 | 3.0 | 1.7 |
| TYR | 3.3 | 2.3 | 2.9 | 1.4 |
| TRP | 3.1 | 4.3 | 3.9 | 2.4 |
| HIS | 2.2 | 2.5 | 2.2 | 1.4 |

**Table S4. Protein-ssDNA complexes used for the calculation of direct hydrogen bonds between non-aromatic amino acids and nucleobases.**

| PDB ID |
|---|
| 1YEG, 1I7D, 1JMC, 1MJE, 1QZH, 1S40, 1ZZI, 2C62, 2CCZ, 2ES2, 2KN7, 2LTT, 2MAP, 2MNA, 2OA8, 2UP1, 2VW9, 2VYE, 3A5U, 3CMU, 3H15, 3ULP, 3VKE, 4GNX, 4GOP, 4HID, 4HIK, 4HIM, 4HIO, 4HJ5, 4HJ7, 4HJ8, 4HJ9, 4HJA, 4I28, 4I2F, 4J1J, 4LJR, 4OU6, 5D8F, 5EAX, 5FHD, 5H1B, 5KEG, 5ODL, 5ORQ, 5TD5, 5USB, 6BUX, 6FWR, 6I52 |

**Table S5. The values of strength $\varepsilon_{ij}^{Hydrogen}$ for hydrogen bonding interactions between different nucleobase and non-aromatic $C_\alpha$ bead pairs.**

| | A | T | G | C |
|---|---|---|---|---|
| GLY | 0.30 | 0.75 | 0.21 | 0.60 |
| ALA | 0.06 | 0.45 | 0.04 | 0.35 |
| VAL | 0.04 | 0.15 | 0.26 | 0.30 |
| LEU | 0.01 | 0.10 | 0.04 | 0.10 |
| ILE | 0.04 | 0.10 | 0.36 | 0.10 |
| SER | 0.30 | 1.50 | 0.61 | 1.00 |
| THR | 0.19 | 0.50 | 0.51 | 0.60 |
| CYS | 0.01 | 0.01 | 0.01 | 0.01 |
| MET | 0.07 | 0.41 | 0.06 | 0.15 |
| PRO | 0.01 | 0.15 | 0.01 | 0.01 |
| ASN | 0.02 | 0.41 | 0.18 | 0.20 |
| GLN | 0.01 | 0.41 | 0.04 | 0.41 |

### Various other Parameters used to Model the Protein and ssDNA:

**Table S6. Mass and Radius of Amino acid Residues.**

| Amino acid (single letter code) | Mass (Da) | Radius of $C_{\alpha}$ atom ($\AA$) |
|---|---|---|
| Isoleucine (I) | 131.1 | 2.0 |
| Lysine (K) | 146.1 | 2.0 |
| Phenylalanine (F) | 165.2 | 2.0 |
| Threonine (T) | 119.1 | 2.0 |
| Tryptophan (W) | 204.2 | 2.0 |
| Valine (V) | 117.1 | 2.0 |
| Arginine (R) | 174.2 | 2.0 |
| Histidine (H) | 155.1 | 2.0 |
| Alanine (A) | 89.0 | 2.0 |
| Asparagine (N) | 132.1 | 2.0 |
| Leucine (L) | 131.1 | 2.0 |
| Methionine (M) | 149.2 | 2.0 |
| AsparticAcid (D) | 133.1 | 2.0 |
| Cytosine (C) | 121.1 | 2.0 |
| Glutamic acid (E) | 147.1 | 2.0 |
| Glutamine (Q) | 146.1 | 2.0 |
| Glysine (G) | 75.0 | 2.0 |
| Proline (P) | 115.1 | 2.0 |
| Serine (S) | 105.0 | 2.0 |
| Tyrosine (Y) | 181.1 | 2.0 |

**Table S7. Mass and Radius for Nucleic acid components.**

| | Mass (amu) | Radius ($\AA$) |
|---|---|---|
| Phosphate | 94.97 | 2.25 |
| Sugar | 83.11 | 3.20 |
| Adenine (A) | 134.1 | 2.70 |
| Thymine (T) | 125.1 | 3.55 |
| Guanine (G) | 150.1 | 2.45 |
| Cytosine (C) | 110.1 | 3.20 |

**Table S8. Equilibrium bond lengths ($r^0$) for the CG DNA. P(5′) and S(5′) denotes the phosphate and sugar in the 5′direction of the adjacent site whereas the P(3′) and S(3′) denotes the phosphate and sugar in the 3′ direction.**

| Bond | $r^0$($\AA$) |
|---|---|
| P (5′) - S | 3.899 |
| S - P (3′) | 3.559 |
| S - A | 4.670 |
| S - T | 4.189 |
| S - G | 4.829 |
| S - C | 3.844 |

**Table S9. Equilibrium bend angles ($\theta^0$) for the CG DNA.**

| Bend | $\theta^0$(°) |
|---|---|
| S - P - S | 94.49 |
| P - S - P | 120.15 |
| P - S - A | 103.53 |
| P - S - T | 92.06 |
| P - S - G | 107.40 |
| P - S - C | 103.79 |
| A - S - P | 112.07 |
| T - S - P | 116.68 |
| G - S - P | 110.12 |
| C - S - P | 110.33 |

**Table S10. Equilibrium dihedral angles ($\varphi^0$) for the CG DNA. (5′)P and (5′)S denotes the phosphate and sugar in the 5′direction of the adjacent site whereas the P(3′) and S(3′) denotes the phosphate and sugar in the 3′ direction.**

| Dihedrals | $\varphi^0$(°) | $\sigma_\varphi$ |
|---|---|---|
| (5′)P - S - P - S(3′) | -154.79 | 0.30 |
| (5′)S - P - S - P(3′) | -179.17 | 0.30 |

**Table S11. Details of the Studied Protein-ssDNA Systems.**

| PDB ID | Sequence of ssDNA | # Nucleotides | pH | Temperature |
|---|---|---|---|---|
| 4GNX | poly T | 25 | NULL | NULL |
| 6I52 | poly T | 20 | 7.4 | NULL |
| 2LTT | poly T | 17 | 7.5 | 298K |
| 2CCZ | poly T | 15 | 6.5 | 100K |
| 4OU6 | poly T | 10 | NULL | NULL |
| 5ODL | poly T | 9 | 7.0 | 293K |
| 2ES2 | poly T | 6 | 6.5 | 293K |
| 2MNA | poly T | 6 | NULL | NULL |
| 2C62 | TTTTTTTTTTTTTTTTTTTG | 20 | 6.75 | 100K |
| 5ZG9 | TTTTTTTTTTTTTTTTTTTG | 20 | 7.5 | 291K |
| 1EYG | poly C | 35 | 7.5 | 291K |
| 1JMC | poly C | 8 | 6.8 | 100K |
| 1S40 | GTGTGGGTGTG | 11 | 7.0 | 303K |
| 2UP1 | TAGGGTTAGGG | 11 | 8.5 | 100K |
| 3VKE | ACCCCA | 6 | 4.2 | 293K |
| 1QZH | GGTTAC | 6 | 8.5 | 289K |
| 1ZZI | CTCCCC | 6 | 8.5 | 294K |
| 2KN7 | CAGTGGCTGA | 10 | 5.2 | 293.8K |
| 2MAP | TGTCAAA | 7 | 6.5 | 303K |
| 4HIO | GGTAACGGT | 9 | 7.0 | 290K |
| 4HJA | ACGGTTACGGT | 11 | 7.0 | 290K |
| 5ORQ | TTAGGGTTAG | 10 | 4.6 | 298K |
| 5USB | GGTTACGGT | 9 | 8.0 | 277K |
| 6BUX | AATCCCAAA | 9 | 5.0 | 277K |
| 6KBS | CGGTCGATTC | 10 | 4.0 | 293K |

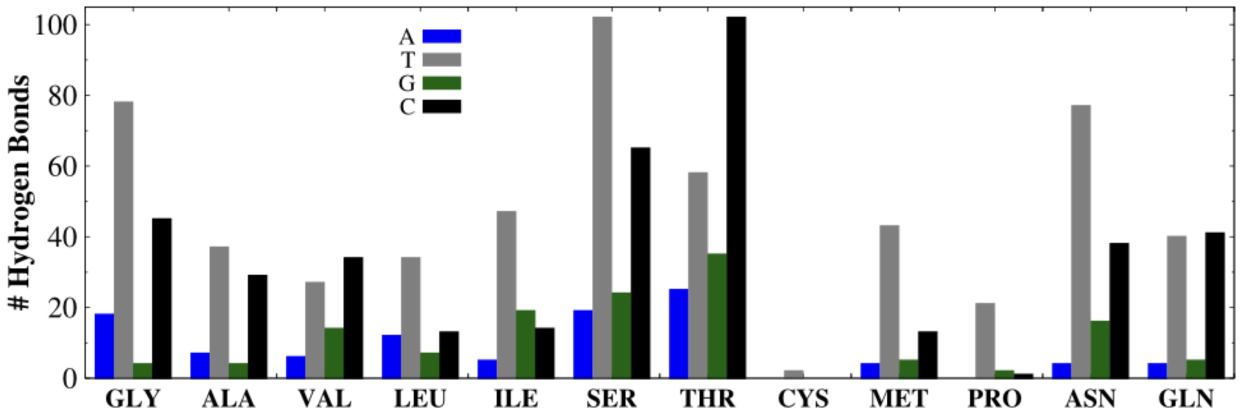


**Figure S1.** The number of direct hydrogen bonds between non-aromatic amino acids and nucleobases derived for 51 protein-ssDNA complexes (see Table S4) from *Hydrogen Bond Computing Server, HBCS (19)*.

**Binding of poly T sequences**

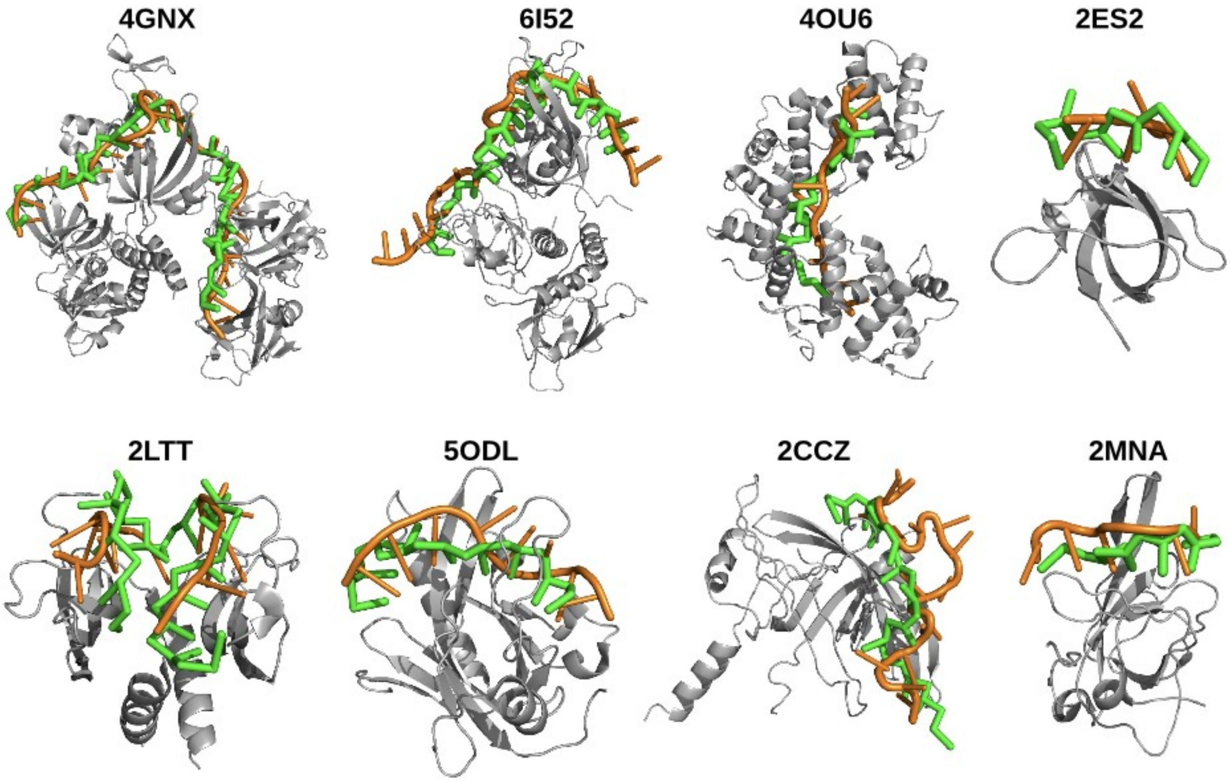


**Binding of poly C sequences**

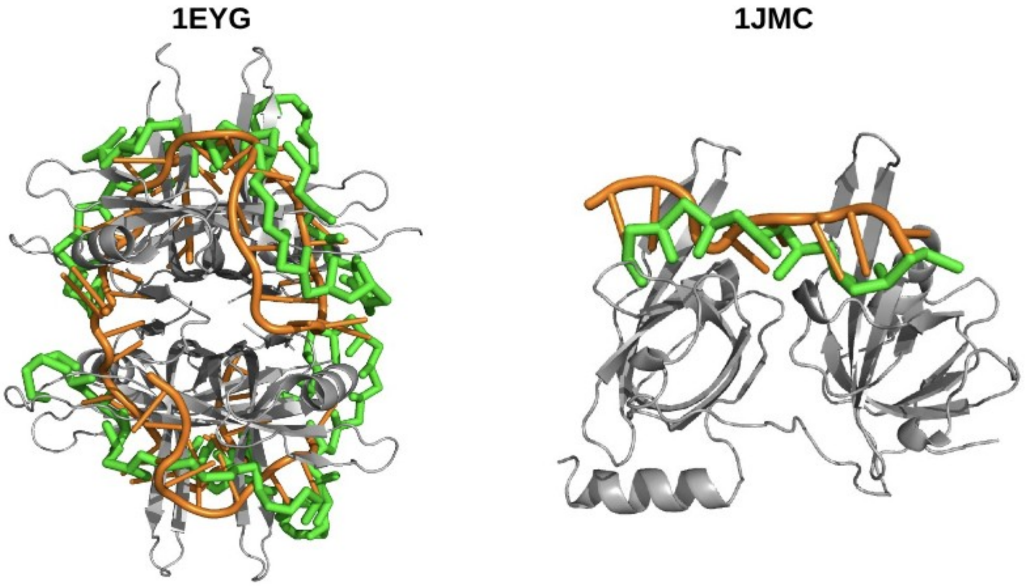

**Binding of mixed sequences**

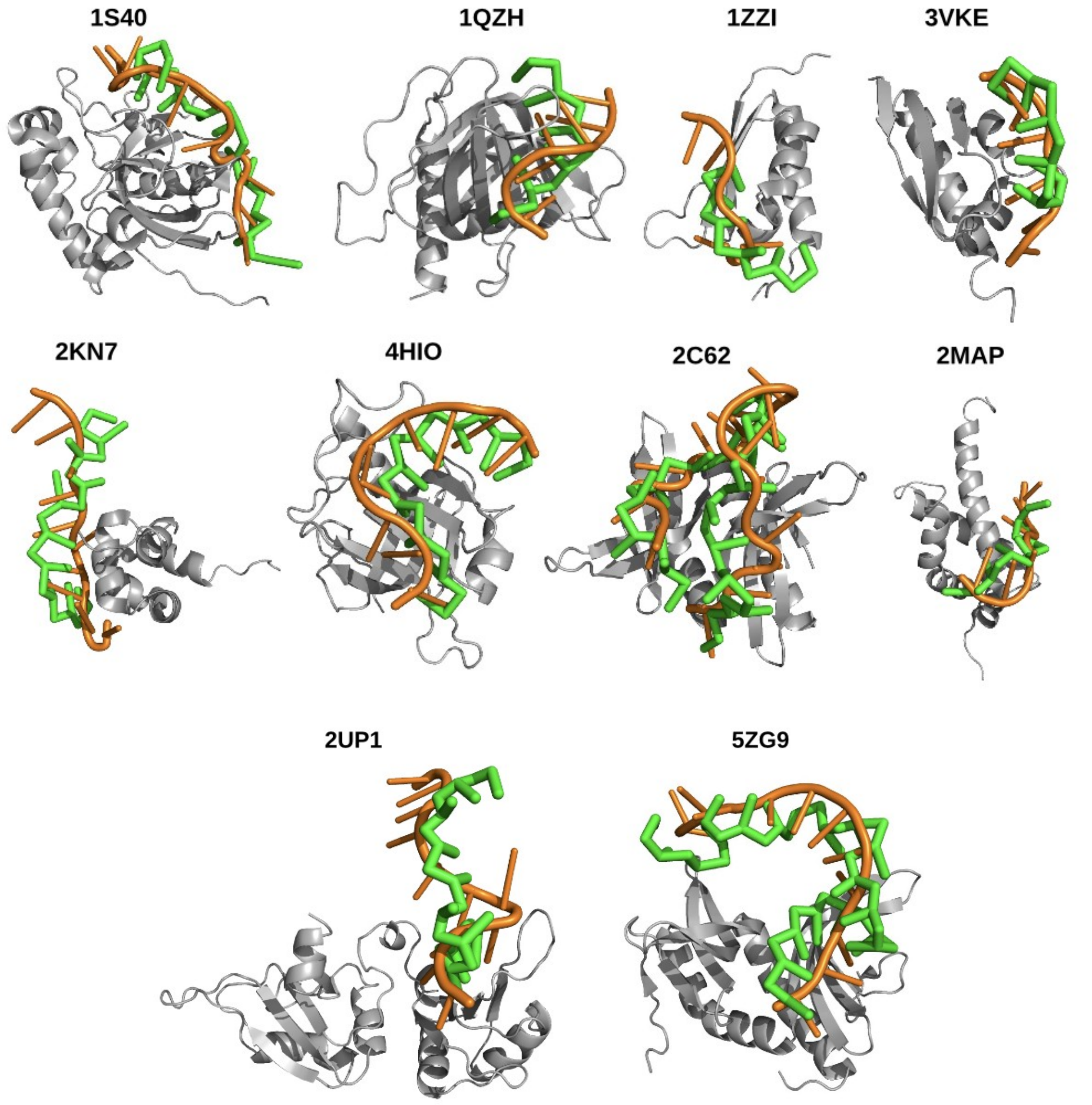


**Figure S2.** Structures corresponding to the proper bound state conformation obtained from our coarse-grained simulations are shown for various protein-ssDNA complexes that differ in terms of the length of the ssDNA and the sequence composition of the nucleotides. For comparison, the simulated ssDNA (green color) after achieving the bound conformation is superimposed with the ssDNA in the corresponding crystal structure (orange color). The proteins are shown in cartoon representation, grey color.

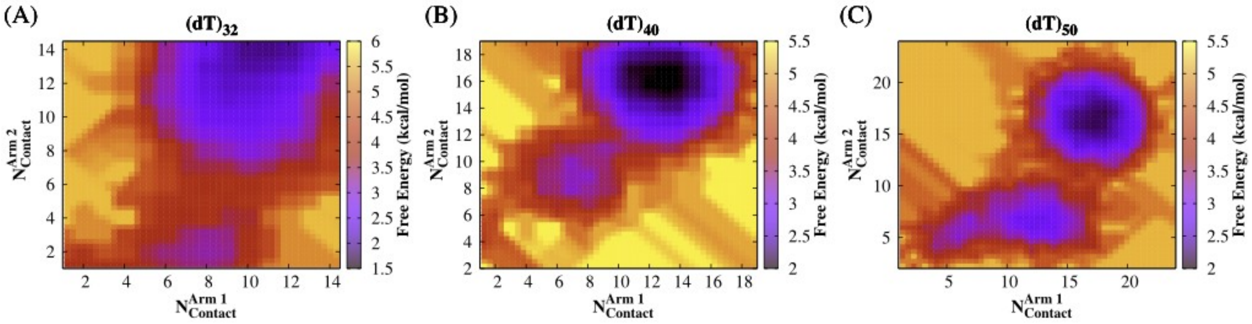


**Figure S3.** Two-dimensional free energy profiles for $(dT)_{32}$, $(dT)_{40}$and $(dT)_{50}$ length of ssDNA as a function of arm-wise contacts ($N_{contact}^{Arm\,1}$and $N_{contact}^{Arm\,2}$) between nucleobases and the closest amino acid residues (within a distance cutoff of 6 Å).

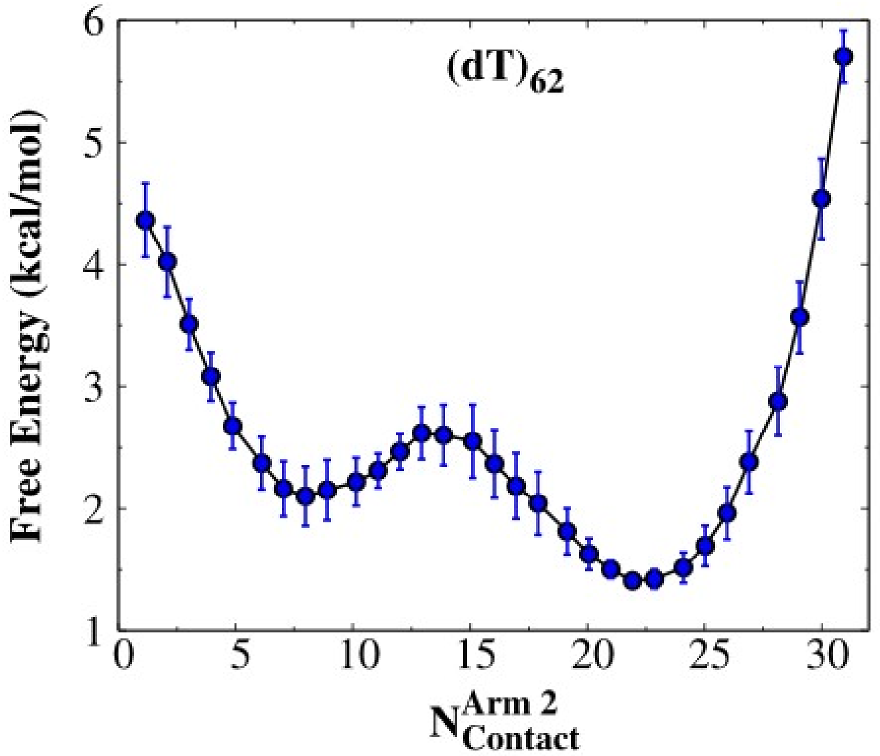


**Figure S4.** Free energy for long ($(dT)_{62}$) length of ssDNA as a function of number of contacts in *arm-2* ($N_{contact}^{Arm\,2}$) between nucleobases and the closest amino acid residues (within a distance cutoff of 6 Å).

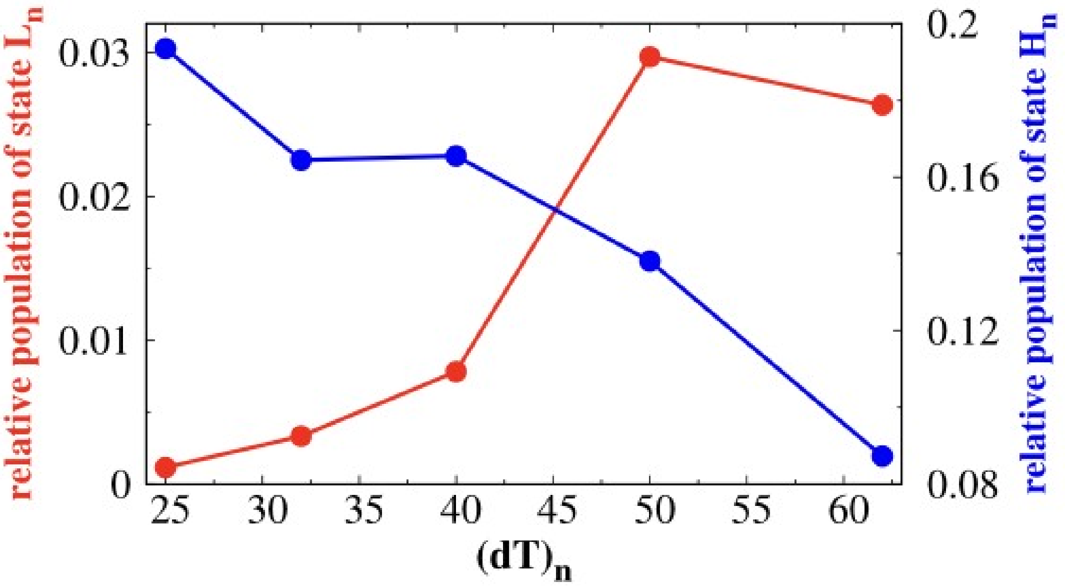


**Figure S5.** The relative population of states $L_n$ (red color) and $H_n$ (blue color) as a function of $(dT)_n$, where ‘n’ indicates the length of the ssDNA. $L_n$ corresponds to a linear configuration of ssDNA perpendicular to the RPA surface and $H_n$ represents the bound state with a topology resembling a ‘horse-shoe’ shaped bent structure.

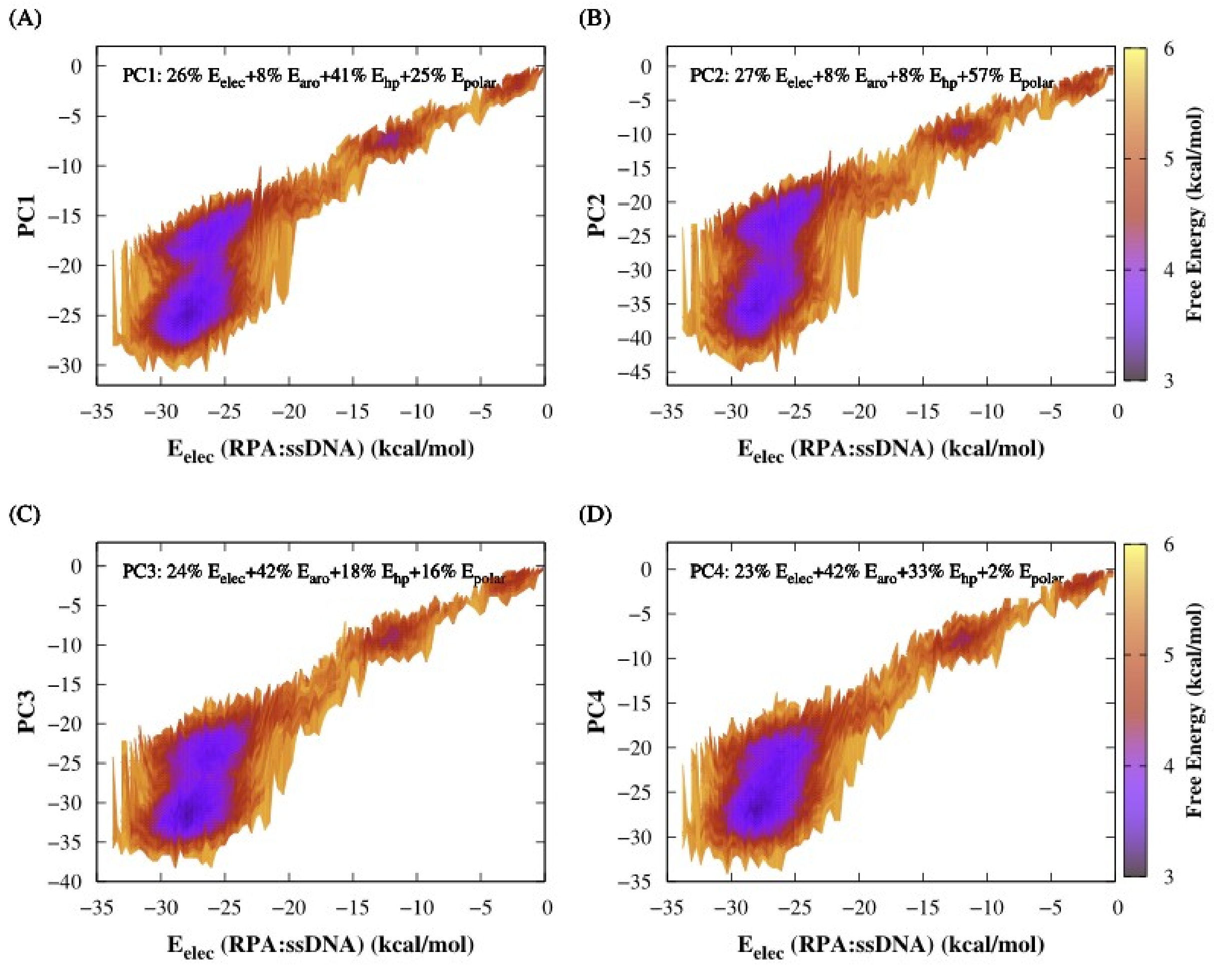


**Figure S6.** Two-dimensional free energy surfaces for binding of RPA with $(dT)_{32}$as a function of various principal components (PCs). Each panel (A-D) shows the binding free energy is projected on a particular PC (PC1 to PC4) along with a common coordinate $E_{elec}$(RPA:ssDNA). The number percentage in each PC represents the fraction of the collective basis variables (see Table S12 for details).

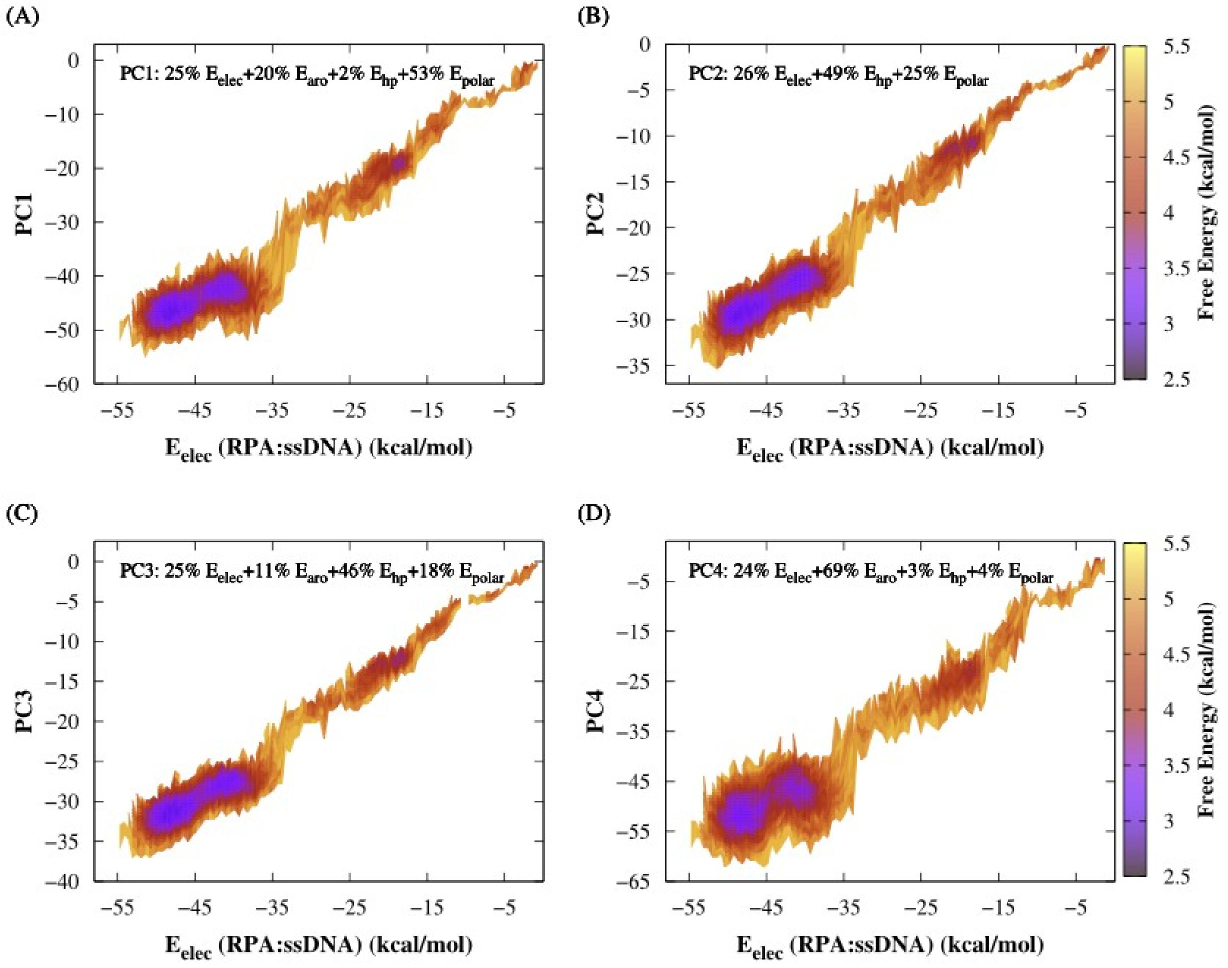


**Figure S7.** Two-dimensional free energy surfaces for binding of RPA with $(dT)_{50}$ as a function of various principal components (PCs). Each panel (A-D) shows the binding free energy is projected on a particular PC (PC1 to PC4) along with a common coordinate $E_{elec}$ (RPA:ssDNA). The number percentage in each PC represents the fraction of the collective basis variables (see Table S13 for details).

**Table S12. Coefficients for Principal Components (PCs) of $(dT)_{32}$ binding mode**

| | $E_{elec}$ | $E_{aro}$ | $E_{hp}$ | $E_{polar}$ |
|---|---|---|---|---|
| **PC1 (48.0%)[a]** | **0.262[b]** | **0.080** | **0.412** | **0.246** |
| **PC2 (35.6%)** | **0.270** | **0.078** | **0.084** | **0.568** |
| **PC3 (12.1%)** | **0.236** | **0.417** | **0.177** | **0.164** |
| **PC4 (4.3%)** | **0.232** | **0.424** | **0.327** | **0.017** |

**Table S13. Coefficients for Principal Components (PCs) of $(dT)_{50}$ binding mode**

| | $E_{elec}$ | $E_{aro}$ | $E_{hp}$ | $E_{polar}$ |
|---|---|---|---|---|
| **PC1 (39.0%)[a]** | **0.250[b]** | **0.203** | **0.021** | **0.526** |
| **PC2 (29.0%)** | **0.257** | **0** | **0.491** | **0.252** |
| **PC3 (29.0%)** | **0.254** | **0.113** | **0.455** | **0.177** |
| **PC4 (3.0%)** | **0.240** | **0.685** | **0.032** | **0.044** |

$E_{elec}$: electrostatic energy between RPA and ssDNA, $E_{aro}$: aromatic interaction energy between aromatic amino acids and nucleobases, $E_{hp}$: the interaction energy between hydrophobic amino acids and nucleobases, $E_{polar}$: the interaction energy between ssDNA base atoms and polar amino beads.

[a]The percentage of total variance described by each PC. [b]The values obtained from the square of the principal component coefficients represent the fraction of the input collective basis variables.

These values are used as coefficients for the linear combination in each PC. Note that $\sum_{i=1}^{4} b_i = 1$

where i is the index of the collective basis variable.

In RPA-$(dT)_{32}$ binding, the first three PCs (PC1 to PC3) account for 95.7% of the total variance. In these, the contributions from electrostatic, aromatic, hydrophobic and polar interactions in RPA-(dT)62 binding are 26.17%, 12.19%, 26.02% and 35.54% respectively. The percentages are estimated by the formula:

$$Percentage\ of\ E_k = \frac{\sum_{i=PC1}^{PC3} \left[ E_k(i)\ x\ percentage\ of\ total\ variance\ explained\ by\ (i) \right]}{\sum_{i=PC1}^{PC3} percentage\ of\ total\ variance\ explained\ by\ (i)}\ x\ 100\,\%$$

where $k = elec, aro, hp, polar$

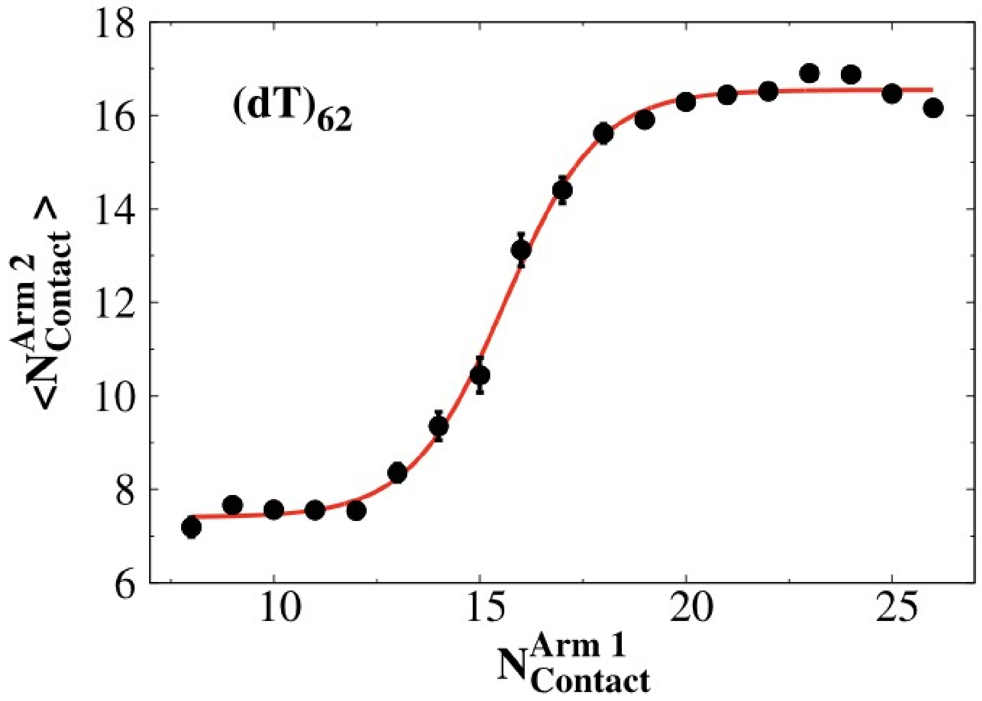


**Figure S8.** Cooperative binding of $(dT)_{62}$ length of ssDNA with RPA. The average number of contacts in *arm-2* ($N_{contact}^{Arm\,2}$) between nucleobases and closest protein residues is plotted with respect to the number of contacts in *arm-1* ($N_{contact}^{Arm\,1}$). The red curve corresponds to the best fit sigmoidal curve.

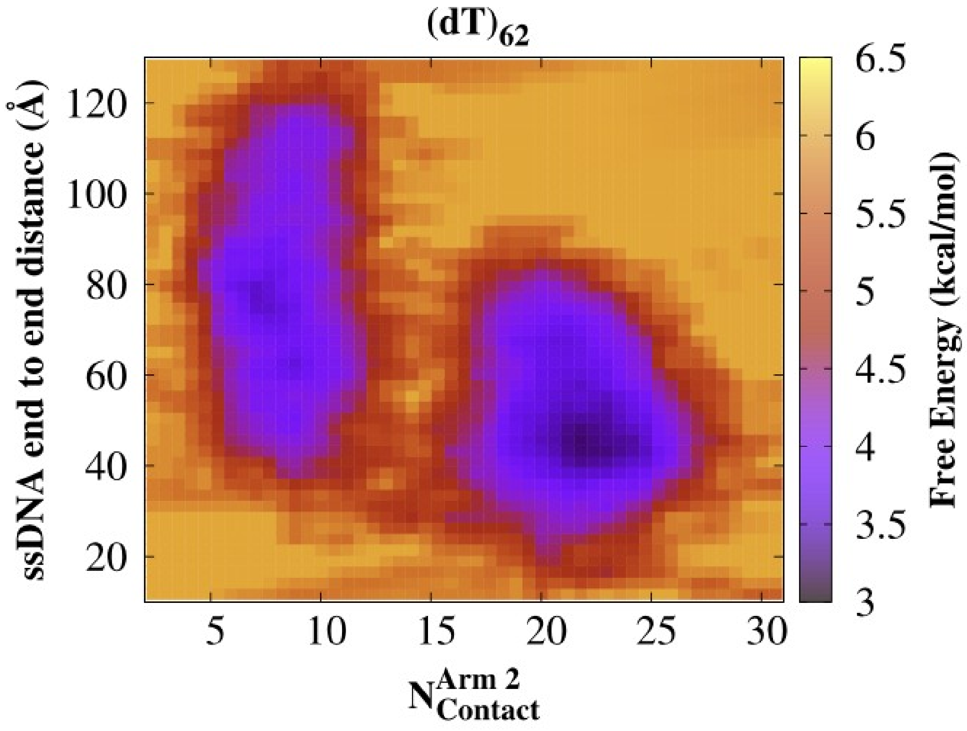


**Figure S9.** Two dimensional free energy surface for long ($(dT)_{62}$) length of ssDNA as a function of ssDNA end-to-end distance and the number of contacts in arm-2 ($N_{contact}^{Arm\,2}$) only.

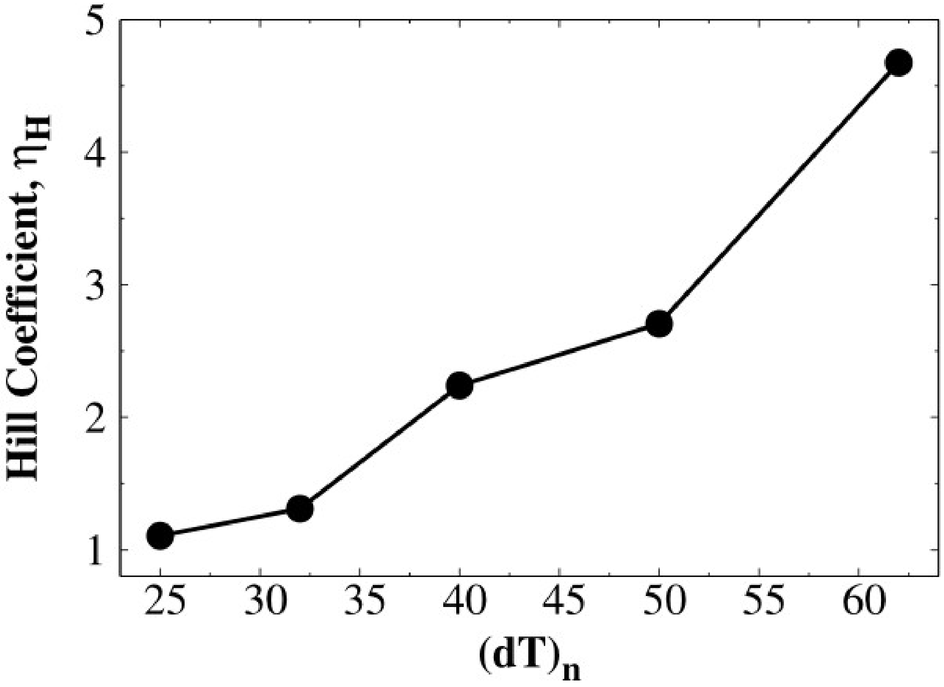


**Figure S10.** Hill coefficient ($\eta_H$) as function of $(dT)_n$, where 'n' indicates the length of the ssDNA. The value of the Hill coefficient describes the cooperativity of the RPA binding with ssDNA. $\eta_H>1$ corresponds to the positive cooperative binding.

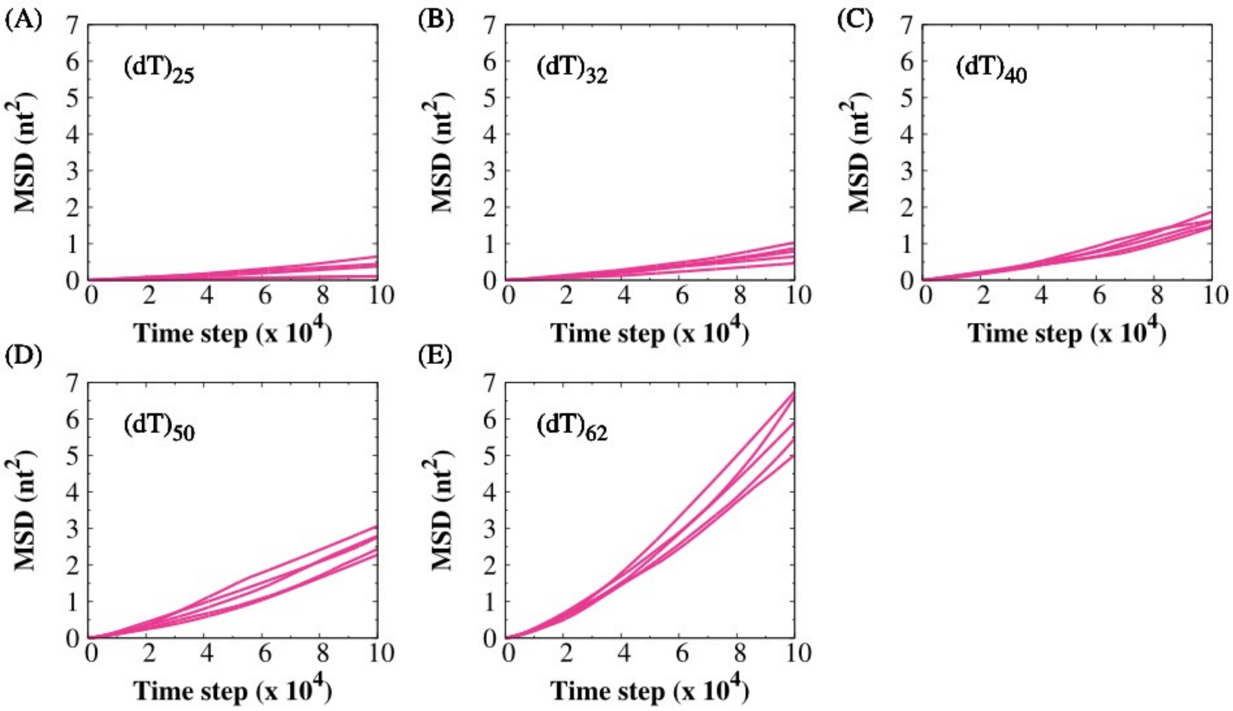


**Figure S11.** The mean square displacement (MSD) of nucleotide bases at site F236 of DBD-A as a function of time. The raw MSD data for $(dT)_{25}$, $(dT)_{32}$, $(dT)_{40}$, $(dT)_{50}$ and $(dT)_{62}$ are shown in each panel.

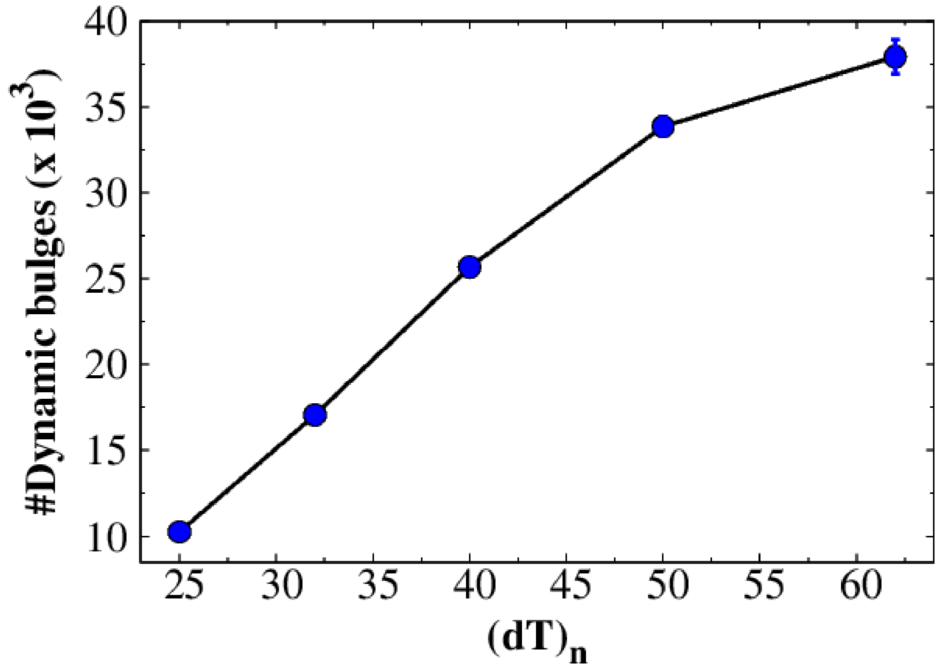


**Figure S12.** The number of 'dynamic bulges' as a function of of $(dT)_n$, where 'n' indicates the length of the ssDNA.

### References.

12. K. A. Wilson, J. L. Kellie, S. D. Wetmore, DNA–protein π-interactions in nature: abundance, structure, composition and strength of contacts between aromatic amino acids and DNA nucleobases or deoxyribose sugar. *Nucleic Acids Research* **42**, 6726–6741

(2014).

13. G. Mishra, Y. Levy, Molecular determinants of the interactions between proteins and ssDNA. *Proc. Natl. Acad. Sci. U. S. A.* **112**, 5033–5038 (2015).

14. J. Fan, N. P. Pavletich, Structure and conformational change of a replication protein A heterotrimer bound to ssDNA. *Genes Dev.* **26**, 2337–2347 (2012).

15. E. Jeong, H. Kim, S.-W. Lee, K. Han, Discovering the interaction propensities of amino acids and nucleotides from protein-RNA complexes. *Mol. Cells* **16**, 161–167 (2003).

16. A. Marcovitz, Y. Levy, Chapter 10. Sliding Dynamics Along DNA: A Molecular Perspective. *RSC Biomolecular Sciences*, 236–262.

17. T. Veitshans, D. Klimov, D. Thirumalai, Protein folding kinetics: timescales, pathways and energy landscapes in terms of sequence-dependent properties. *Folding and Design* **2**, 1–22 (1997).

18. A. Pal, Y. Levy, Structure, stability and specificity of the binding of ssDNA and ssRNA with proteins. *PLoS Comput. Biol.* **15**, e1006768 (2019).

19. M. Gurusaran, *et al.*, Hydrogen Bonds Computing Server (HBCS): an online web server to compute hydrogen-bond interactions and their precision. *Journal of Applied Crystallography* **49**, 642–645 (2016).
